# Molecular insights into dynamic protein structures by high-contrast crosslinking mass spectrometry

**DOI:** 10.1101/2024.09.02.610668

**Authors:** Zhuo Angel Chen, Eva Absmeier, James Stowell, Ludwig Roman Sinn, Shabih Shakeel, Tamara Sijacki, Kendra Njo, Kolja Stahl, Edward Rullmann, Francis J. O’Reilly, Lori A. Passmore, Juri Rappsilber

## Abstract

Proteins are comprised of structured domains and dynamic regions, and both are essential for biological function. However, studying dynamic regions is challenging using most structural biology methods, including crosslinking mass spectrometry. Here, we dramatically improve the usefulness of distance restraints from crosslinking MS by taking advantage of short-lived reactive species generated from diazirine-based photo-crosslinking. This leads to a clear view of complex topologies and conformational changes, including in dynamic regions. We demonstrate that photo-crosslinking MS data can be used to model flexible regions and conformational changes in the DNA repair complexes; Fanconi Anemia core complex and FANCD2-FANCI. In addition, we obtain new insights into the architecture and arrangement of the highly flexible CCR4-NOT mRNA deadenylation complex. The improved contrast of photo-crosslinking empowers structural biology by providing clearer structural insights into dynamic biological systems that have eluded other structural biology approaches.

## Introduction

Protein dynamics play pivotal roles in shaping protein functions. This includes structural flexibility, conformational changes and rearrangements within individual proteins and multi-protein complexes. Additionally, approximately 40% of eukaryotic proteins contain substantial intrinsically disordered regions^1^. Dynamic flexibilty present challenges to structural biological methods designed mostly to study structured (rigid) protein domains. While crosslinking mass spectrometry (MS) has emerged as a transformative tool in structural biology^2–4^, the data produced represents protein dynamics as a composite of states, making it challenging to unravel the intricacies of dynamic processes.

Crosslinking MS involves the creation of new covalent bonds between nearby residues within a protein or protein complex^5^. These artificially created bonds are then detected using proteomics, which reveals the specific residues that are linked together. This information sheds light on the structure of the protein or complex in solution, even within a complicated sample such as organelles and whole cells^6–8^. Protein crosslinking requires special reactive chemicals. The most commonly used crosslinkers usually contain two terminal NHS esters, which primarily facilitate the linking of nearby lysines, but also serine, threonine, and tyrosine residues. Frequently used examples of crosslinkers are bissulfosuccinimidyl suberate (BS3) and bis(2,5-dioxopyrrolidin-1-yl) 3,3′-sulfinyldipropionate (DSSO)^9^. For crosslinking, the NHS ester based crosslinkers are typically incubated with protein samples for tens of minutes^10,11^. When one end of crosslinker reacts with a protein residue, the other side remains reactive for minutes to hours before it hydrolyses (Extended Data Fig. 1a, b)^12^. Thus, under such conditions, these crosslinkers may trap conformational states of proteins that are rarely occupied in solution and amplify their presence in the ensemble, leading to many crosslinks in flexible regions. Also, crosslinking may trap chance encounters between nearby molecules. There is no easy way to filter out crosslinks representing rare conformations or chance encounters, which makes the structural information difficult to interpret.

An alternative, though less commonly employed, group of chemicals for exploring protein structure is photo-crosslinkers, particularly succinimidyl 4,4′-azipentanoate (SDA). Similar to BS3 and DSSO, SDA features an NHS ester, but only on one terminus. On the other end, SDA contains a photoactivatable diazirine (Extended Data Fig. 1c). Upon exposure to UV light (350-365 nm), diazirines react with all amino acids^13^ (Extended Data Fig. 1d), although, at least under some conditions, they favour acidic side chains^14^. This broad reactivity leads to an increase in the structural coverage and the density of crosslinking data, however, it also complicates the analysis process, as it disperses the mass spectrometric signal among more products and enlarges the database search space. Despite these challenges, recent improvements in data acquisition and database searches have made it possible to use photo-crosslinking MS for studying the structure of large protein complexes^15,16^ and even entire proteomes^17^. The length of SDA bridges is considerably shorter (3.9 Å) than that of other crosslinkers (BS3: 11.4 Å), falling within the range of interactions seen in hydrogen bonding (2.7-3.3 Å^18^) and hydrophobic contacts (3.8-9.5 Å^19^). The limited length of SDA may also be more suitable for investigations into contact sites^15^. Most importantly, the activated diazirine has an extremely brief lifetime (∼2 ns^20^) and is therefore most likely to reveal conformational states that are abundant in solution^21^. As a result, it provides structural information primarily on the most prevalent conformer, or group of conformers, in the sample.

Here, we aim to explore the possibility that the homo-bifunctional NHS ester crosslinkers most frequently employed in crosslinking MS investigations, including BS3 and DSSO, generally have an adverse impact on the contrast, i.e., the ability to distinguish between specific interactions and random contacts. This limits the structural interpretation of the resulting data, particularly in the case of dynamic and flexible systems. We also show that the use of photo-crosslinkers substantially improves contrast in crosslinking MS, likely because of short crosslinker lifetimes. This provides an exciting new tool to investigate the structure of flexible protein complexes and their dynamic behaviour.

## Results

### Photo-crosslinking defines the interface of a dynamic subunit within a structured complex

The megadalton FA core complex contains at least eight core subunits (FANCA, B, C, E, F, G, L, FAAP100) and is essential for DNA repair and genomic stability. We previously determined a structure of the chicken FA core complex using cryo-EM and crosslinking MS with BS3^22,23^. Our model did not include FANCA which could not be confidently modelled into the cryoEM maps, presumably due to flexibility. In particular, many BS3 crosslinks were found between FANCA and all other subunits and so these could not be used to assign the location of FANCA in the complex (Fig. 1a). In a subsequent cryoEM study, FANCA was modelled in the human FA core complex where it was present in two copies and was stabilised by an additional subunit, FAAP20^24^.

**Figure 1:**
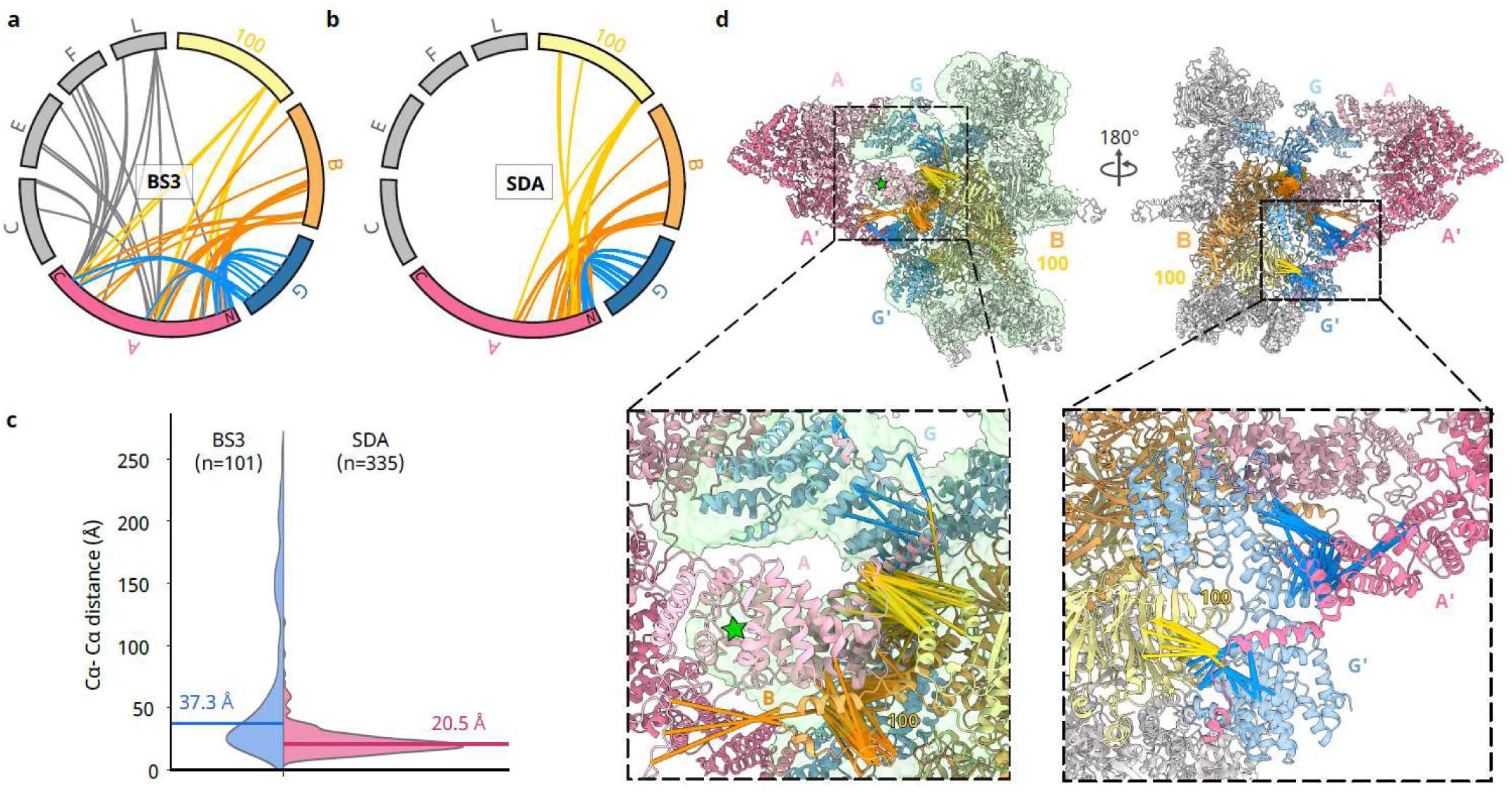
SDA crosslinking reveals the location of the FANCA subunit in the chicken FA core complex. a. 112 BS3 crosslinked residue pairs identified between FANCA and other subunits of the FA core complex. Each crosslink is colour-coded according to the specific subunit FANCA was crosslinked to. b. 343 SDA crosslinked residue pairs identified between FANCA and other subunits of the FA core complex. Crosslinks are colour-coded as shown in (A). c. Violin plot illustrating the distribution of C_α_-C_α_ distances between residue pairs crosslinked by BS3 (blue) and SDA (pink), as measured in a homology model of the chicken FA core complex. The homology model was generated based on the 3D structure model of the human FA core complex (PDB|7KZP). The median C_α_-C_α_ distance of 37.4 Å for the BS3 crosslinks and 20.4 Å for the SDA crosslinks is indicated by solid lines. d. Visualization of SDA crosslinks between FANCA and other subunits in the homology model of the chicken FA core complex. The model is displayed in cartoon mode, with the front and back views shown. Crosslinks are represented as solid lines and colour-coded as described in (A). In the front view (left), the homology model is shown within the cryo-EM map of the chicken FA core complex (EMD-10290). Two regions in the atomic model showed a high concentration of crosslinks, which are highlighted in the amplified views. In the left amplified view, the green star indicates unexplained density within the cryo-EM map of the chicken FA core complex, which coincides with the N-terminal region of one FANCA in the homology model.

We wondered if the shorter reactive lifetime of photo-crosslinkers would produce distance restraints that indicate how FANCA binds to the rest of the FA core subunits in our chicken FA core complex. We therefore crosslinked the FA core complex with the sulfo-derivative of SDA (referred to as SDA throughout the text) instead of BS3. Analysis of this complex identified 3,459 SDA crosslinks, compared to 843 with BS3 (Fig. 1, Extended Data Fig. 2, Supplemental data 1). As the SDA and BS3 analyses varied in depth (see Methods), a difference in the number of identified crosslinks is expected. Excluding FANCA, the crosslink-based interaction network of subunits was very similar in both cases, even with the much larger number of inter-subunit crosslinks observed with SDA (541 compared to 122 with BS3) (Extended Data Fig. 2).

FANCA crosslinked to other subunits of the FA core complex via 343 crosslinks with SDA and 112 with BS3. For BS3, FANCA crosslinked to other subunits at three regions along the length of the protein, linking to every other protein in the complex and preventing unambiguous assignment of FANCA to unexplained EM density (Fig. 1a). BS3 and SDA self-crosslinks covered the entire length of FANCA (Extended Data Fig. 3), but SDA heteromeric crosslinks of FANCA exclusively involved its N-terminal region, which crosslinked to FANCB, FANCG, and FAAP100 (Fig. 1b). These subunits are positioned next to the unassigned density in the cryoEM map of the chicken FA core complex (Fig. 1a), suggesting that the N-terminal region of FANCA is located in this area.

To check the agreement of both BS3 and SDA data with the cryoEM map, we built a homology model for the chicken FA core complex using the human cryoEM structure, which included two copies of FANCA (Supplemental data 2). With a linker length of 3.9 Å and assuming two side chains of 5.5 Å being bridged, SDA crosslinks should be no longer than about 15 Å (Cα-Cα distance). The experimental crosslinking length is likely longer than this due to the flexibility of proteins in solution compared to static structural models. Despite the high flexibility of FANCA, SDA crosslinks between FANCA and other FA core complex subunits agree very well with the homology model (Fig. 1c). The median Cα-Cα distance of SDA crosslinked residues is 20.5 Å, with 92% being closer than 35 Å (72% closer than 25 Å). In contrast, the median Cα-Cα distance of BS3 crosslinked residues is 37.3 Å (Fig.1d). Although the crosslinkers differ in linker length (SDA: 3.9 Å, BS3: 11.4 Å) this does not suffice to explain the observed differences in Cα-Cα distances.

The cryo-EM data did not yield conclusive information regarding the stoichiometry of FANCA within the chicken CCR4-NOT complex. Our data show that SDA links the N-terminal region of FANCA to both the N- and C-terminal regions of FAAP100. The thirteen crosslinks to the C-terminus of FAAP100 (C_α_-C_α_ distances of 18.3±4.4 Å) place the N-terminal region of FANCA in the unexplained EM density (Fig. 1d). However, this position results in FANCA being distant from the N-terminus of either of the two copies of FAAP100 (102.1±3.8 Å or 117.7±4.8 Å, 8 crosslinks) (Extended Data Fig. 4). In the homology model, the N-terminal region of the second copy of FANCA is proximal to the C-terminal segment of one FAAP100 (20.7±1.8 Å), consistent with two copies of FANCA in both the human and chicken complexes .

Overall, our analysis of the FA core complex strongly suggests that rapid photo-crosslinking improves the contrast of crosslink data in dynamic regions, providing clear insights into the molecular architecture of protein interfaces. In contrast, the commonly used NHS ester chemistry produces many artifactual crosslinks, which hinder the interpretation of crosslinks as distance restraints for protein (complex) modeling

### Photo-crosslinking reveals alternate conformational states

The distinct behaviour of the two crosslinking approaches for a complex with flexible elements raised the question of how crosslinking chemistry affects the quantitative analysis of a complex with multiple conformational states. FANCD2-FANCI (D2-I) is a DNA clamp that exists in open and closed states, where both states are predominantly structured but a large rotation of one subunit is required to transition between states^25,26^. D2-I closure enables its ubiquitination, which is a key step that activates DNA repair by the FA pathway. Using cryoEM and quantitative crosslinking MS with SDA, we previously showed that D2-I exists in a conformational equilibrium between the two states. Phosphomimetic mutations on FANCI shift this equilibrium towards the closed state, promoting the ubiquitination of the D2-I complex^27^.

To understand how the crosslinker chemistry influences quantitative crosslinking MS analysis, we repeated the crosslinking of the wild-type (D2-I^WT^) and phosphomimetic (D2-I^3D^) complexes with BS3. Unlike for the FA core complex, the analysis here for both crosslinkers started with identical amounts of the D2-I complexes and used the same mass spectrometry analysis methods. BS3 returned 316 quantified crosslinks, and SDA returned 233 quantified crosslinks (Supplemental data 3). Within the respective dataset (SDA or BS3), all quantified crosslinks are present in both D2-I^WT^ and D2-I^3D^ indicating the simultaneous presence of the same conformers in solution.

If the equilibrium between open and closed conformers is different for D2-I^WT^ and D2-I^3D^, we would expect to observe this through differences in the abundance of conformer-specific crosslinks. For individual crosslinks, larger intensity ratios between D2-I^WT^ and D2-I^3D^ are observed for SDA (194 < 2-fold, 39 > 2-fold) than for BS3 (304 < 2-fold, 12 > 2-fold), with a maximal ratio of 30 for SDA compared to 3.8 for BS3 (Fig. 2a, b). This reflects greater differences between conformational states with SDA. Thus, SDA preserves population differences between D2-I^WT^ and D2-I^3D^ better than BS3, which revealed how D2-I’s conformational equilibrium affects its activation^27^. In contrast, the newly acquired BS3 data could not reveal these conformational differences, presumably due to conformational averaging over the longer time-scales of the crosslinking (Fig. 2c). Overall, photo-crosslinking captures the proportion of alternative conformational states of a structured protein complex in solution, likely due to the rapid nature of the crosslinking event.

**Figure 2:**
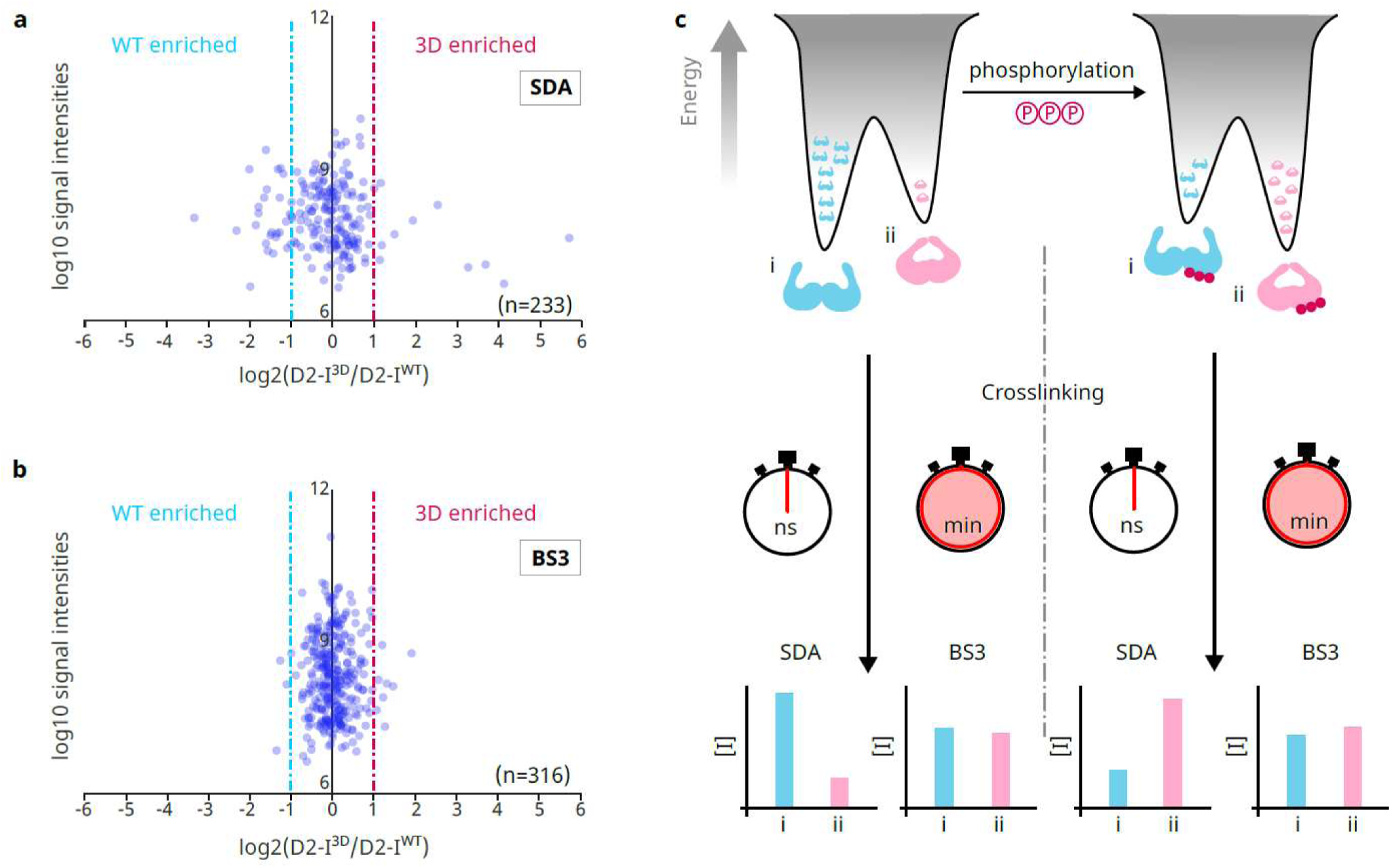
Photo-crosslinking preserves conformational ensembles of the D2-I complex. a. Plot of quantified SDA cross-links, with log2 D2-I^3D^/D2-I^WT^ signal ratios plotted against the log10 intensities of the cross-links. Vertical dashed lines mark the twofold enrichment threshold. b. Plot of quantified BS3 cross-links, with log2 D2-I^3D^/D2-I^WT^ signal ratios plotted against the log10 intensities of the cross-links. Vertical dashed lines mark the twofold enrichment threshold. The chemical structure of BS3 is illustrated. The green pentagon represents the NHS ester-based reaction group. c. A model of the observed difference in the D2-I^WT^ vs. D2-I^3D^ quantitation results using two different types of crosslinkers (as shown in panels A and B). The diazirine-based SDA crosslinking only remains reactive for milliseconds once activated, which better captures the true population of the conformations in the ensemble. NHS ester-based BS3 crosslinking stays reactive in solution for tens of minutes, which can lead to conformational averaging.

### Photo-crosslinking reveals the topology of a dynamic multiprotein complex

Next, we wondered if crosslinking can reveal the topology of a highly dynamic complex which is too flexible for structure determination by cryoEM. For this, we chose the CCR4-NOT complex, which contains six conserved, core subunits and plays crucial roles in various cellular processes, including mRNA degradation, translation repression and transcription regulation^28^. In the human CCR4-NOT complex, CNOT1 acts as a scaffold protein to organize the other subunits into four functional modules, CNOT2-CNOT3, CNOT6/6L-CNOT7/8, CNOT9, CNOT10-CNOT11^29^. While the structures of these modules have been individually characterized through crystallography and cryoEM, the 3D architecture of these modules within the presumably highly dynamic complex remains unclear^30^.

We crosslinked recombinant human CCR4-NOT complex (eight subunits) using BS3 and SDA. We identified 1,284 BS3-crosslinks and 958 SDA-crosslinks (Supplemental data 4). All known subunit-subunit interactions are captured in the BS3 data. In addition, however, every subunit is crosslinked to every other CCR4-NOT subunit, including all 28 theoretically possible subunit pairs of the complex, preventing the assignment of a clear subunit topology (Fig. 3a). In contrast, the SDA data report proximity for 11 of the 28 theoretically possible subunit pairs, including 8 of the 9 known interactions. Specifically, for crosslinks within individual subunits in available crystal structures, both BS3 and SDA bridge residues in proximity, with median Cɑ-Cɑ distances of 15.4 Å for BS3 crosslinks and 13.7 Å for SDA crosslinks. For crosslinks between subunits, a similar distance is observed for SDA crosslinked residues (14.2 Å) but BS3 resulted in many more long-distance crosslinks, with a median Cɑ-Cɑ distance of 34.7 Å (Fig. 3b).

**Figure 3:**
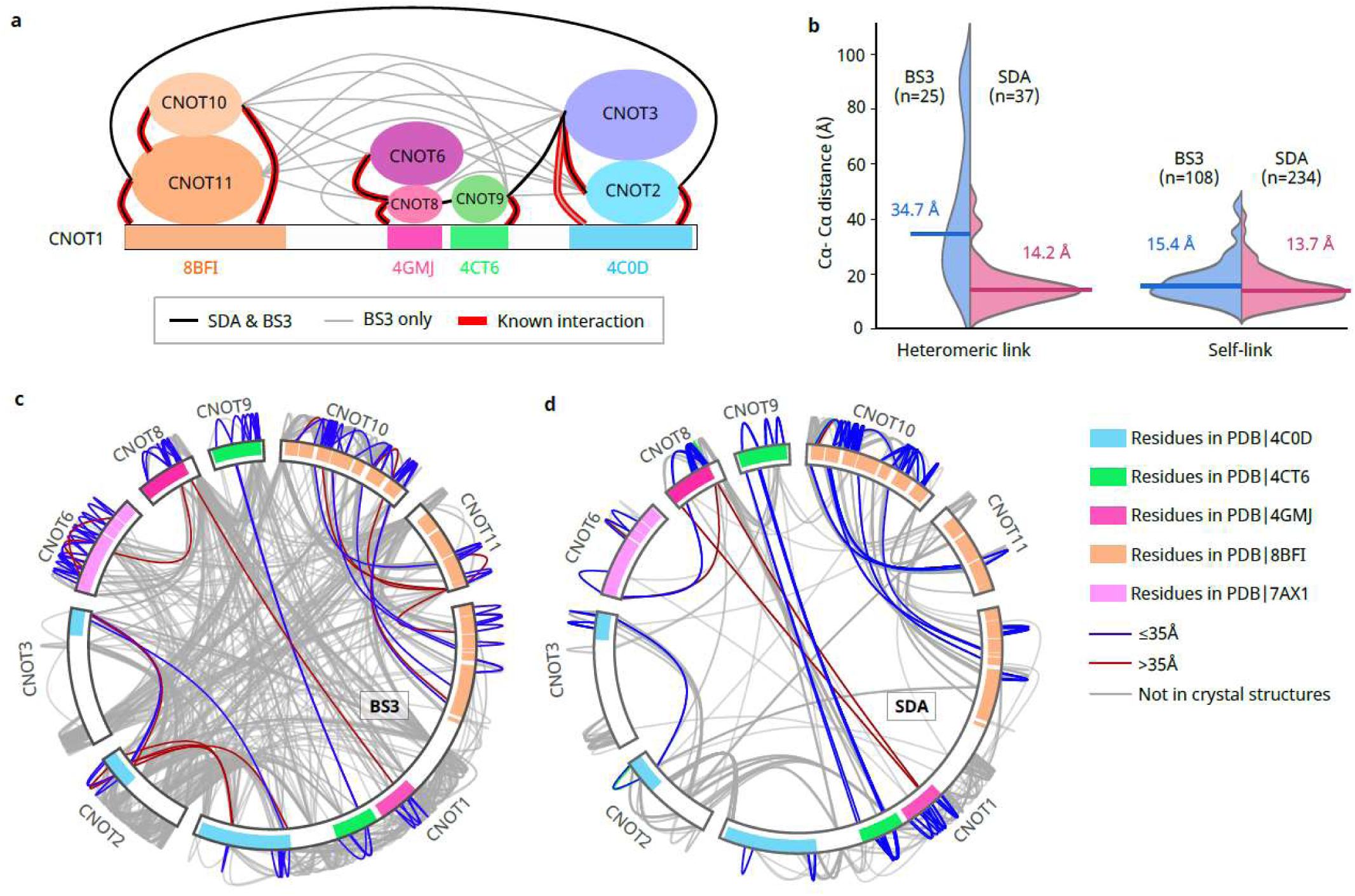
Photo-crosslinking reveals the topology of the CCR4-NOT complex. a. The crosslinking network of subunits within the CCR4-NOT complex. In the schematic representation of the CCR4-NOT complex, CNOT1 is shown as a bar, while other subunits are shown as ovals, scaled to the size of the proteins. The ovals are positioned on CNOT1 where they interact, as revealed by crystallographic data. Thick red lines depict interactions between subunits as defined in the crystal structures. Segments of CNOT1 that interact with other subunits in the four functional modules are highlighted with coloured shades: CNOT1^1–685^ (orange), CNOT1^1093–1317^ (pink), CNOT1^1356–1588^ (green) and CNOT1^1847–2361^ (blue). The corresponding crystal structures revealing these interactions are also indicated with PDB IDs. Black and grey lines show crosslinks between subunits with a minimum of three unique residue pairs: black lines indicate crosslinks observed with both SDA and BS3, while grey lines indicate those observed only with BS3. BS3 crosslinks were observed between every pair of subunits in the complex. b. Violin plot illustrating the distribution of Cα-Cα distances between residue pairs crosslinked by BS3 (blue) and SDA (pink), measured in the crystal structures of the functional modules of the human CCR4-NOT complex: CNOT1-CNOT2-CNOT3 (PDB 4C0D), CNOT1-CNOT8-CNOT6 (PDB 4GMJ, 7AX1), CNOT1-CNOT9 (PDB 4CT6), and CNOT1-CNOT10-CNOT11 (PDB 8BFI). The median Cα-Cα distances of BS3 and SDA crosslinks are indicated by solid lines for both self-links and heteromeric links. c. A map of BS3 crosslinks in the CCR4-NOT complex. Crosslinks that can be measured in the crystal structures of the functional modules (as described in panel A) are highlighted in blue when the Cα-Cα distances between the linked residues are less than or equal to 35 Å, and in red when the distances are greater than 35 Å. The segments of the subunits present in the crystal structures are shaded with corresponding colours. d. A map of SDA crosslinks identified in the CCR4-NOT complex. Crosslinks measurable in the crystal structures of the functional modules (as described in panel A) are coloured as in panel C. The segments of the subunits present in the crystal structures are shaded with corresponding colours.

To rationalise this stark contrast, we investigated the crosslinks of the scaffold protein CNOT1. BS3 crosslinks fail to define clear interaction regions between other subunits and the 2,376-residue CNOT1 protein, even in cases where interfaces are already known from X-ray crystallography of complex fragments. (Fig. 3c, Extended Data Fig. 5a). BS3 data therefore suggest a highly dynamic arrangement of the complex at the level of which proteins interact and how they interact. In contrast, the known interfaces are clearly reflected by the SDA crosslinks (Fig. 3d, Extended Data Fig. 5b). In effect, only SDA recapitulated the known contact regions between the subunits.

In addition to known interactions, three new crosslinked subunit pairs were identified connecting the functional modules: CNOT9-CNOT3, CNOT9-CNOT8, and CNOT2-CNOT11. CNOT9 and CNOT3 are part of two functional modules that have neighbouring interaction regions on CNOT1. This also applies to CNOT9 and CNOT8. In addition, CNOT9 and CNOT8 overlap in their crosslinks to CNOT1, further supporting their proximity within the complex. In contrast, CNOT2 and CNOT11 are from two functional modules that interact with CNOT1 at the N-terminal and the C-terminal regions respectively. These crosslinks suggest the spatial proximity between the two functional modules in the complex.

Taken together, photo-crosslinking recapitulated the known aspects of CCR4-NOT complex topology and provided new insights into subunit arrangement, whereas data obtained with a homobifunctional NHS ester crosslinker were inconclusive. Interestingly, all the crosslinks that were observed between established functional modules involve Intrinsically disordered regions (IDRs).

### Photo-crosslinking reveals IDR interactions in the CCR4-NOT complex

CCR4-NOT contains multiple subunits with IDRs that may contribute to the dynamic nature of the complex (Supplemental data 5). A substantial fraction of our crosslinks involves IDRs (631 or 50% of 1,284 BS3 crosslinks and 321 or 33% of 958 SDA crosslinks). Notably, 95% of the SDA IDR crosslinks connect to structured segments, providing insights into how the IDRs might be anchored to the complex. This was also the case for 75% of the BS3 IDR crosslinks.

For the short IDR in CNOT11 (residues 285-320), the BS3 and SDA data are in good agreement (Extended Data Fig. 6a,b). In contrast, for less constrained IDRs, BS3 and SDA give different results. For example, at the termini of proteins (the N-terminal IDR of CNOT11 and the C-terminal IDR of CNOT10, Extended Data Fig. 6) or within IDRs with longer sequences (Fig. 4a, b, c; Extended Data Fig. 7), BS3 crosslinks are found to every other CCR4-NOT subunit. This is likely indicative of a highly dynamic IDR and may not reflect that these subunits are normally in close proximity. SDA crosslinks within these regions are more defined. BS3 also provides minimal information regarding the structural arrangement of the CNOT2 IDR, one of the longest IDRs in the complex (Fig. 4a). Intriguingly, with SDA CNOT2^37–51^ and CNOT2^153–167^ crosslink with each other and to adjacent regions of the structured C-terminal domain of CNOT2 (Fig. 4a), providing insight into the structural arrangement of CNOT2.

**Figure 4:**
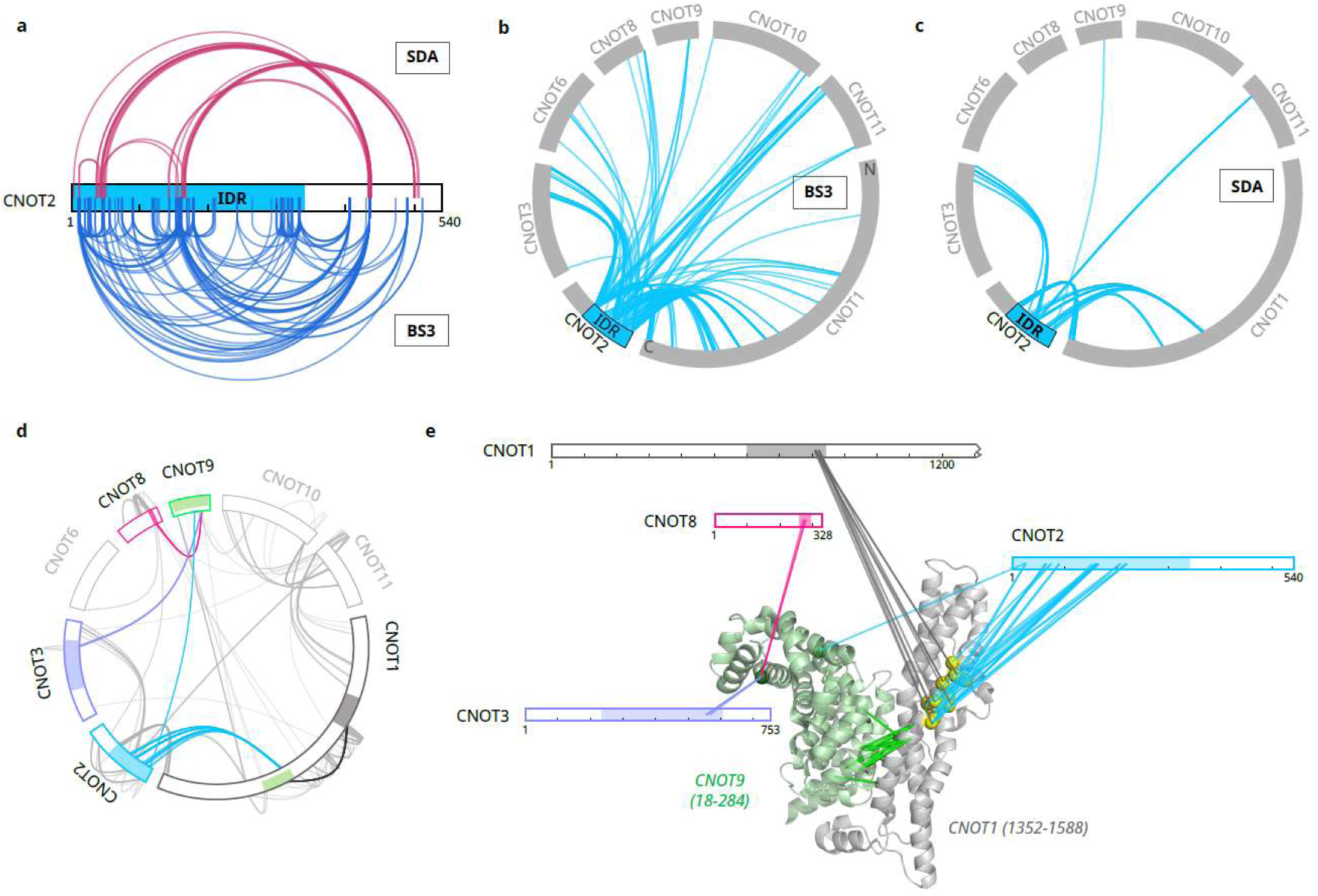
Photo-crosslinking provides insight into the structural arrangement of IDRs in the CCR4-NOT complex. a. Self-links from the IDR in CNOT2: CNOT2 is represented as a bar with the IDR indicated in cyan. SDA crosslinks are depicted as arches in dark pink, and BS3 crosslinks as arches in blue. b. BS3 crosslinks from CNOT2 IDR (cyan) to other subunits of the CCR4-NOT complex. c. SDA crosslinks from CNOT2 IDR (cyan) to other subunits of the CCR4-NOT complex. d. The CNOT1-CNOT9 interacting region (green; PDB 4CT6) appears as a crosslinking hub for IDRs in CNOT1 (dark grey), CNOT2 (cyan), CNOT3 (slate), and CNOT8 (pink). Crosslinks from the highlighted IDRs to the crosslinking hub are coloured accordingly. Other subunits and IDR crosslinks are shown in light grey on the crosslinking map. e. Crosslinking sites of IDRs depicted on the crystal structure of the CNOT1-CNOT9 interacting regions (cartoon; PDB 4CT6). CNOT1 (1-1300), CNOT2, CNOT3, and CNOT8 are depicted as bars with IDRs indicated in colours as in panel C. Crosslinks are shown as lines connecting residues in IDRs to C-α atoms of crosslinked residues in the crystal structure (spheres). An IDR crosslinking hub in CNOT1 is highlighted in yellow. Crosslinks between CNOT1 and CNOT9 are depicted in green lines on the structure.

Distinct contacts are also evident from SDA crosslinks within the NOT module (CNOT1^1847–2361^– CNOT2-CNOT3) (Extended Data Fig. 8): the C-terminal region of the CNOT2^IDR^ exclusively crosslinked to the C-terminal region of CNOT1 and a short stretch (307-333) in the centre of the CNOT2^IDR^ crosslinked to the C-terminal region of CNOT3 (Extended Data Fig. 8). Ten IDR crosslinks from the SDA data fall within reported crystal structures, with Cα-Cα distances of 15.1±2.9 Å and IDRs that are in sequence close to crystallised regions display plausible crosslink footprints (Extended Data Fig. 6c).

We also found a number of interactions between the CNOT2^IDR^ and the CNOT1^1356–1588^–CNOT9 module (Fig. 4d,e). CNOT9 is a highly conserved subunit in the CCR4-NOT complex, with its concave surface reported as a binding site for the unstructured regions of several RNA-binding proteins^31–36^. Our data showed that IDRs from CNOT3 and CNOT8 also crosslinked to CNOT9 within this concave surface (Fig. 4d,e). Unexpectedly, two groups of residues from both the N-terminal part of CNOT2^IDR^ (25-90) and the centre of CNOT2^IDR^ (153-219) were crosslinked to CNOT1^1400–1410^, which is adjacent to the CNOT1-CNOT9 interface (Fig. 4d,e). CNOT1^1400–1410^ also crosslinked to two distinct stretches in IDRs that are similar in length but different in composition: one in CNOT1 itself (808-822: KLGTSGLNQPTFQQS) and one in the C-terminal purification tag of CNOT8 (310-328: SAGSAAGSGAGWSHPQFEK). These crosslinks suggest that within the human CCR4-NOT complex, the CNOT1^1356–1588^–CNOT9 module also serves as a binding hub for IDRs.

The associations of CNOT1^1400–1410^ with multiple IDRs and the absence of crosslinks between these IDRs suggest a network of alternative interactions. We made mutations in CNOT1^1400–1410^ to disrupt the binding of IDRs, but these did not affect *in vitro* complex assembly, recruitment of CCR4-NOT to RNA or nuclease activity of the complex (Extended Data Fig. 8). Instead, the CNOT1 IDR binding site may influence the recruitment of regulatory factors in cells. For example, the RNA-binding TTP protein binds to two regions in the CCR4-NOT complex, CNOT9 and CNOT1^800–999^ ^37^. The recruitment of the CNOT1^808–822^ IDR to CNOT1^1400–1410^ brings both regions together, possibly assembling a composite binding site.

Clusters of SDA crosslinks from the centre of CNOT2^IDR^ (153-242) span an apparent network of contacts to structured regions of several functional modules, including the C-termini of CNOT2 and CNOT3 in the NOT module, both CNOT1 and CNOT9 in the CNOT1-CNOT9 module, CNOT11 in the CNOT10-CNOT11 module and the CNOT1 MIF4G-C domain (Extended Data Fig. 9). These contacts between IDRs and structured domains may contribute to the orchestration of CCR4-NOT function. The N-terminal and central parts of CNOT2^IDR^ are highly enriched for phosphorylation sites^38,39^, with over 48% of all phosphorylation sites in the CCR4-NOT complex located in the region. This long IDR region in CNOT2 may therefore serve a regulatory role in the CCR4-NOT complex.

The residue-level SDA data revealed a number of interesting details regarding subunit interactions of the CCR4-NOT complex that include IDRs. Our data show that the improved contrast of photo-crosslinking extends to IDRs, providing evidence for interactions of IDRs mostly with structured regions that now define starting points for investigating the in-cell regulation of the complex.

## Discussion

NHS ester crosslinkers are routinely used for analysing the structure of isolated or reconstituted protein complexes, and increasingly also in complex mixtures. Here, we demonstrate limitations of these reagents in flexible and dynamic complexes. Because of solvation effects and higher susceptibility to proteinaceous nucleophiles, NHS esters preferentially target proteins over water^12^, remaining active for long periods (minutes to hours). Homobifunctional NHS ester crosslinkering can therefore accumulate through chance or rare encounters in dynamic protein systems. In contrast, a protein adorned with a photo-activatable group reacts within nanoseconds after exposure to UV light^20^, either with a bound entity, with adjacent residues in the sequence or with water. This short lifetime bestows high contrast to photo-crosslinking, unveiling remarkably meaningful interactions in our study (Fig. 2c). The combination of a short lifetime and broad reactivity makes diazirines also imortant reagents for proximity labelling, especially for highly dynamic proteins^40^.

The limitations of homobifunctional NHS ester crosslinkers may extend to crowded environments. For example, studies of *in situ* protein interactions often unveil high percentage of unexpected biological links^6–8,41^. Only some of these have been confirmed, such as the existence of a novel proteinaceous pyruvate dehydrogenase (PDH) inhibitor in *Bacillus subtilis*^6,8^. It is possible that some of the in-situ crosslinks are spurious due to the long timescales required for crosslinking.

In the future, development of reagents that increase the number of residues that can be crosslinked and digested would expand the capabilities and coverage of crosslinking MS. While the diazarine of SDA can crosslink many if not all residues, it requires a K, S, T, or Y residue on the NHS ester side. Likewise, trypsin being used as protease requires K or R for cleavage, albeit alternative proteases can be employed in crosslinking MS^42,43^. As can be seen here in the IDR of CNOT3, which lacks tryptic cleavage sites, these residue restrictions may lead to absence of data (Extended Data Fig. 10) and point towards future development directions of the technology.

We establish photo-crosslinking as a high-contrast chemical procedure that, when paired with mass spectrometry, is capable of revealing the topology of highly dynamic complexes, conformational rearrangements, and interaction interfaces involving IDRs. Its ability to capture elusive structural features may make photo-crosslinking a valuable tool also in combination with other analytical approaches, beyond the mass spectrometric detection of linkage sites

## Author’s contribution

S.S. prepared and crosslinked the FA core complex. F.O.R. and L.R.S. performed CLMS analysis on the crosslinked FA core complex. T.S. prepared and crosslinked the D2-I complexes, while Z.A.C. conducted CLMS analysis on these crosslinked D2-I complexes. E.A. prepared and crosslinked the CCR4-NOT complex, with K.N. carrying out the CLMS analysis. Z.A.C. processed and analyzed all data. E.R. generated the homology model of the chicken FA core complex. K.S. identified the IDRs in the CCR4-NOT complex. J.S. prepared the mutant CCR4-NOT complex and performed differential scanning fluorimetry and in vitro deadenylation assays on both the wild-type and mutant CCR4-NOT complexes. Z.A.C. and J.R. wrote the manuscript with input from all authors.

## Acknowledgements

We thank Andrea Graziadei for discussions and J.G. Shi (LMB Baculovirus facility) for assistance. This work was funded by the Deutsche Forschungsgemeinschaft (DFG, German Research Foundation) under Germanýs Excellence Strategy – EXC 2008 – 390540038 – UniSysCat and the Wellcome Trust through a Discovery Award (227434) and core funding for the Wellcome Centre for Cell Biology (203149); by the MRC, as part of United Kingdom Research and Innovation (also known as UK Research and Innovation), MRC file reference number MC_U105192715 (L.A.P.); the European Union’s Horizon 2020 research and innovation program (European Research Council Consolidator grant agreement no. 725685) (to L.A.P.) and a postdoctoral fellowship of the Deutsche Forschungsgemeinschaft (the German Research Foundation, project no. 429892960) to E.A.

## Data availability

The mass spectrometry proteomics data have been deposited to the ProteomeXchange Consortium via the PRIDE^44^ partner repository with the dataset identifier PXD054492 and 10.6019/PXD054492.

Reviewer account details:

Username

Password: rsczmkZkbl0p

## Methods

### Preparation of protein complexes

The *G. gallus* FA core complex was expressed and purified from baculovirus-mediated insect cell expression, following an established protocol^45^. Briefly, Sf9 cells were infected with the recombinant baculovirus carrying the FA core complex genes (FANCA, FANCG, FANCB, FANCL, FAAP100, FANCC, FANCE and FANCF) with a StrepII tag on FANCC. The expressed FA core complex was isolated and purified using affinity, heparin and anion-exchange chromatography. The purified FA core complex was resuspended in a buffer containing 50 mM HEPES at pH 8.0, approximately 500 mM NaCl, and 1 mM TCEP.

The expression and purification of *G. gallus* FANCD2 and two FANCI constructs (FANCI^WT^ and FANCI^3D^) were carried out as previously described^27^. In brief, the Sf9 cells were transfected with the recombinant baculovirus carrying full-length FANCD2 gene that contains a C-terminal extension with a 3C protease site and double StrepII tag or full-length *FANCI* gene that contains a C-terminal extension with a 6x His tag respectively. The FANCI^3D^ construct was generated by inserting a 130-bp sequence (gBlocks, IDT) encoding S558D, S561D and T567D mutations into the linearised pACEBac FANCI^WT^. Cells were harvested at 90% viability (typically 3–4 days post infection). After lysing the cells and clarifying the lysate by ultracentrifugation, the expressed FANCD2 or FANCI proteins were isolated and purified using affinity, heparin and size exclusion chromatography. To prepare D2-I^WT^ and D2-I^3D^, 3.15 μM of FANCD2 was incubated with either 3.15 μM FANCI^WT^ or 3.15 μM FANCI^3D^ in assay buffer (50 mM HEPES pH 8.0, 150 mM NaCl, 1 mM TCEP) at 4 °C for 30 min before crosslinking reaction.

Human CCR4-NOT complex was expressed with a BigBac approach in insect cells and purified as described^46^. Briefly, 6 l SF9 cell culture was infected with V0 virus and CCR4-NOT complex was expressed with a low multiplicity of infection strategy for 72-84 h. Cells were harvested by centrifugation at 800g, flash frozen in liquid nitrogen and stored at -80°C. Cells were resuspended in 250 ml lysis buffer (150 mM NaCl, 50 mM HEPES-NaOH pH 7.5, 2 mM magnesium acetate, 0.1 mM TCEP, 0.1% v/v NP-40), supplemented with EDTA-free protease inhibitor tablets and 3 mg DNaseI and lysed by shearing through a needle. Cell lysate was cleared by centrifugation at 200,000 g for 25 min at 4°C and filtered supernatant was incubated with 3 ml Biolock solution, before batch-binding to 10 ml bed-volume equilibrated Strep-resin for 1h in the cold room. Subsequently, beads were separated from the flow-through by gravity flow column, washed four times with wash buffer (150 mM NaCl, 40 mM HEPES-NaOH pH 7.5, 2 mM magnesium acetate, 0.1 mM TCEP) and protein was eluted by adding five times 10 ml elution buffer (150 mM NaCl, 40 mM HEPES-NaOH pH 7.5, 2 mM magnesium acetate, 0.1 mM TCEP, 5 mM Desthiobiotin). Protein-containing fractions were pooled and diluted with 40 mM HEPES-NaOh pH 7.5 to a final salt concentration of 80 mM NaCl. Next, the complex was loaded on a 5 ml HiTrap Q HP column, equilibrated in buffer A (80 mM NaCl, 40 mM HEPES-NaOH pH 7.5, 2 mM magnesium acetate, 0.1 mM TCEP) and eluted with a 12 CV linear gradient to 100% buffer B (1 M NaCl, 40 mM HEPES-NaOH pH 7.5, 2 mM magnesium acetate, 0.1 mM TCEP). Fractions which contained stoichiometric complex were pooled and diluted to a final NaCl concentration of 80 mM. Subsequently, the complex was loaded on a 1-ml ResourceQ column equilibrated in buffer A and eluted in a steep gradient (10 CV to 70% buffer B) to concentrate the complex. The complex containing fractions were pooled, flash-frozen in liquid nitrogen and stored at -80°C until usage. Complex concentration was measured using absorbance at 280 nm (A0.1%=0.785).

### BS3 Crosslinking

The purified FA core complex (in 50 mM HEPES pH 8.0, ∼500 mM NaCl and 1 mM TCEP) at a concentration of 7.6 μM was crosslinked with a 100-fold molar ratio of disulfosuccinimidyl suberate (BS3) for 2 h on ice and the reaction was quenched with 50 mM NH_4_HCO_3_ for 30 min at room temperature. The crosslinked samples were then precipitated using cold acetone. The precipitated proteins were then pelleted by centrifugation.

D2-I complex was crosslinked in crosslinking buffer (50 mM HEPES pH 8.0, 150 mM NaCl, 1 mM TCEP) with BS3, with a 1:8 D2-I to BS3 mass ratio. The crosslinking reaction was incubated at 4 °C for 30 minutes and was quenched with NH_4_HCO_3_ at 4 °C for 30 minutes. The crosslinking reaction was carried out in triplicate. The crosslinking products were separated on a 3–8% NuPAGE Tris-acetate gel (Invitrogen), and gel bands corresponding to the cross-linked D2-I heterodimer were excised.

100 µg of purified CCR4-NOT complex was diluted to a final concentration of 0.255 µM in crosslinking buffer (100 mM NaCl, 20 mM HEPES NaOH pH 7.5, 0.1 mM TCEP, 2 mM magnesium acetate). Addition of CCR4-NOT complex added 30 mM NaCl, which resulted in a final salt concentration of 130 mM NaCl. BS3 crosslinker solution was added to a final concentration of 0.05 mM and the sample was incubated for 30 min on ice, before quenching with 100 mM ammonium bicarbonate. The crosslinked complex was then precipitated using cold acetone. The precipitated proteins were then pelleted by centrifugation.

### SDA crosslinking

To keep the crosslinking chemistry identical to BS3 on the NHS ester end, sulfo-SDA (sulfosuccinimidyl 4,4’-azipentanoate), which is a derivative of SDA with improved water solubility ^13^ was used for crosslinking for all protein complexes in this work. The purified FA core complex at a concentration of 8 µM was subjected to crosslinking with different molar ratios of sulfo-SDA. The crosslinking reactions were performed by mixing the FA core complex with sulfo-SDA at 100, 250, 500, 1000, and 2000-fold molar excess ratios. The mixtures were incubated for 2 hours on ice in the dark to enable the reaction of the NHS ester side of the cross-linker to the protein complex. After the incubation, each crosslinking reaction was exposed to UV light at 356 nm for 20 minutes to photoactivate the diazirine group of the cross-linker. The crosslinked samples were then precipitated using cold acetone. The precipitated proteins were then pelleted by centrifugation.

D2-I complex in crosslinking buffer (50 mM HEPES pH 8.0, 150 mM NaCl, 1 mM TCEP) was incubated with sulfo-SDA with a 1:1 protein to crosslinker mass ratio at 4 °C for 30 min. To activate the diazirine group in sulfo-SDA, the sample was irradiated with a UV lamp at 356 nm on ice for 20 min. The crosslinking reaction was conducted in triplicate. The crosslinking products were separated on a 3–8% NuPAGE Tris-acetate gel and gel bands corresponding to the cross-linked D2-I heterodimer were excised.

100 µg of purified CCR4-NOT complex was diluted to a final concentration of 0.255 µM in crosslinking buffer (100 mM NaCl, 20 mM HEPES NaOH pH 7.5, 0.1 mM TCEP, 2 mM magnesium acetate). Addition of CCR4-NOT complex added 30 mM NaCl, which resulted in a final salt concentration of 130 mM NaCl. sulfo-SDA crosslinker solution was added to a final concentration of 0.125 mM and the sample was incubated for 20 min on ice. Next, the sample was irradiated with a UV lamp at 365 nm for 20 min on ice. Subsequently, the reaction was quenched with 100 mM NH_4_HCO_3_. The crosslinked complex was then precipitated using cold acetone and then pelleted by centrifugation.

### Protein digestion

The acetone-precipitated crosslinked FA core complex samples and crosslinked CCR4-NOT samples were in-solution digested as previously described^45^. In brief, the protein pellet was resuspended in 8 M urea prepared in 100 mM NH_4_HCO_3_. Peptides were reduced with 10 mM DTT and alkylated with 50 mM iodoacetamide. Following alkylation, proteins were digested with Lys-C (Pierce) at an enzyme-to-substrate ratio of 1:100 for 4 hours at 22 °C. After diluting the urea to 1.5 M with 100 mM NH_4_HCO_3_ solution, proteins were further digested with trypsin (Pierce) for ca. 15 hours at 22 °C at an enzyme-to-substrate ratio of 1:20. The resulting peptides were then desalted using C18 StageTips ^47^. Peptides from sulfo-SDA crosslinked FA core complex samples with different protein-to-crosslinker ratios were combined before desalting.

The gel bands of the crosslinked D2-I heterodimers were in-gel digested as previously described^27^. Briefly, gel bands were cut into 1 mm cubes, washed and de-stained using 50 mM NH_4_HCO_3_ and acetonitrile. The disulfide bonds were reduced with 10 mM DTT at 37 °C for 30 min and the resulting free thiol groups (-SH) were alkylated with 55 mM iodoacetamide at 23 °C for 20 min in the dark. Subsequently, proteins were digested with trypsin in 45 mM ammonium bicarbonate and 10% acetonitrile with a protein-to-trypsin mass ratio of 50:1. The digestion was incubated at 37 °C for 15 h. The resulting peptides were extracted from the gel pieces and desalted using C18 StageTips^47^.

### Enrichment of crosslinked peptides

Strong cation exchange chromatography was applied to the sulfo-SDA crosslinked and BS3 crosslinked FA core complex to enrich for crosslinked peptides. De-salted peptides were dissolved in mobile phase A (30% acetonitrile (v/v), 10 mM KH2PO4, pH 3) before strong cation exchange chromatography (100 × 2.1 mm Poly Sulfoethyl A column; Poly LC). The separation of the digest used a nonlinear gradient (chicken FA ref 62) into mobile phase B (30% acetonitrile (v/v), 10 mM KH2PO4, pH 3, 1 M KCl) at a flow rate of 200 μl min−1. Ten 1-min fractions in the high-salt range were collected and desalted by StageTips for subsequent liquid chromatography with tandem mass spectrometry (LC-MS/MS) analysis.

Peptide size exclusion chromatography (pepSEC) was applied to the BS3 crosslinked FA core complex, BS3 and sulfo-SDA crosslinked human CCR4-NOT complex to enrich for crosslinked peptides. Peptides from BS3 crosslinked FA core complex were fractionated using a Superdex Peptide 3.2/300 column (GE Healthcare) at a flow rate of 10 μl min−1 using 30% (v/v) acetonitrile and 0.1% (v/v) trifluoroacetic acid as mobile phase. Five 50-μl fractions were collected. Peptides from crosslinked CCR4-NOT complex were fractionated using a Superdex™ 30 Increase 3.2/300 column (GE Healthcare) at a flow rate of 10 μl min−1 using 30% (v/v) acetonitrile and 0.1% (v/v) trifluoroacetic acid as mobile phase. Six 50-μl fractions were collected. The solvent was removed from the collected pepSEC fractions using a vacuum concentrator, and then analyzed by LC-MS/MS.

### LC-MS/MS analysis

LC-MS/MS analysis was carried out using an Orbitrap Fusion Lumos Tribrid mass spectrometer (Thermo Fisher Scientific) coupled online with an Ultimate 3000 RSLCnano system (Dionex, Thermo Fisher Scientific). The sample was separated on a 50 cm EASY-Spray C18 column (Thermo Scientific) operating at 45°C column temperature. Mobile phase A consisted of 0.1% (v/v) formic acid and mobile phase B of 80% v/v acetonitrile with 0.1% v/v formic acid. Samples for analysis were resuspended in 0.1% v/v formic acid, 1.6% v/v acetonitrile. Peptides were loaded and separated at a flow rate of 300 nl min−1.

For the SCX fractions, peptides were separated using a gradient with linear increases from 2% mobile phase B to 12.5% over 10 minutes then to 45% over 80 minutes, followed by an increase to 55% in 2.5 minutes and then a steep increase to 95% mobile phase B in 2.5 min.

Peptides from the pepSEC fractions were also separated with a gradient increase from 2% mobile phase B to 95% mobile phase B over 95 minutes. Similar to the one applied to the SCX fractions, the gradient consists of 4 linear slopes. The endpoints and the steepness of the slopes were optimized for each SEC fraction.

Peptides from the crosslinked D2-I samples were subjected to LC-MS/MS analysis without off-line enrichment for crosslinked peptides. Peptides were separated using a gradient with linear increases first from 2% to 4% B in 1 minute, then from 4% to 40% B in 134 minutes, and finally from 40% to 95% B in 15 minutes

Eluted peptides were ionized by an EASY-Spray source (Thermo Scientific) and introduced directly into the mass spectrometer. The MS data were acquired in data-dependent mode using the top-speed setting with a three-second cycle time. For every cycle, the full scan mass spectrum was recorded in the Orbitrap at a resolution of 120K. Ions with a precursor charge state between 3+ and 7+ were isolated and fragmented employing higher-energy collisional dissociation (HCD). For the crosslinked FA core complex, fragmentation energy was determined by employing a decision tree logic with optimized collision energies ^48^. For crosslinked D2-I samples, a normalized collision energy of 30% was applied. For all crosslinked CCR4-NOT complex samples, stepped collision energies (26%, 28% and 30%) were employed. The fragmentation spectra were then recorded in the Orbitrap with a resolution of 50K for BS3 crosslinked FA core complex, crosslinked D2-I samples, 60K for sulfo-SDA crosslinked FA core complex and all crosslinked CCR4-NOT complex samples. Dynamic exclusion was enabled with a single repeat count and a 60-s exclusion duration.

For each pepSEC fraction of BS3 crosslinked FA core complex, LC-MS/MS acquisition was carried out in duplicate. For each pepSEC fraction of crosslinked CCR4-NOT complex samples, LC-MS/MS acquisition was carried out in triplicate. In a few cases where peptide amounts were limited, acquisitions were only conducted in duplicate.

For the SCX fractions of sulfo-SDA crosslinked FA core complex, nine LC-MS/MS acquisitions were carried out for each fraction. Triplicated acquisitions were performed without the FAIMS device applied, while for the other six acquisitions, the FAIMS device was applied with a different combination of two CV voltages for each acquisition.

### Data processing

For all MS raw data, MS2 peak lists were generated using the MSConvert module in ProteoWizard (version 3.0.11729). Precursor and fragment m/z values were recalibrated. Identification of crosslinked peptides was carried out using xiSEARCH software (https://www.rappsilberlab.org/software/xisearch)^42^. The peak lists derived from different protein complex samples were searched against the protein sequences of corresponding protein subunits. The reversed protein sequences of the protein subunits were used as decoys during the search for error estimation.

For data from the crosslinked FA core complex and the crosslinked CCR4-NOT complex, xiSEARCH version 2.0 was used. The following parameters were used for the searches:

MS accuracy, 3 ppm; MS2 accuracy, 5 ppm; enzyme, trypsin (with full tryptic specificity); allowed number of missed cleavages, 3; missing monoisotopic peak, 2. For all samples, carbamidomethylation on cysteine was set as a fixed modification and oxidation on methionine was set as a variable modification.

For BS3 crosslinked samples, crosslinker was set to BS3. The reaction specificity for BS3 was assumed to be for lysine, serine, threonine, tyrosine and protein N termini. BS3 loop-link, hydrolyzed BS3 on one end and amidated BS3 on one end were set as variable modifications. The maximum variable modification per peptide was set to two.

For sulfo-SDA crosslinked samples, crosslinker was set to SDA. The reaction specificity for SDA was assumed to be for lysine, serine, threonine, tyrosine and protein N termini on the NHS ester end and any amino acids for the diazirine end. SDA loop link and hydrolyzed SDA on the diazirine end were set as variable modifications.

Crosslinked peptide candidates with a minimum of three matched fragment ions (with at least two containing a crosslinked residue) in each crosslinked peptide were filtered using xiFDR (version 2.2.betaB)^49,50^. A false discovery rate of 1% at the residue-pair level was applied, with the ’boost between’ option enabled. Due to the small number of proteins in the sequence database for these isolated protein complexes, PPI level FDR could not be applied. To minimize the likelihood of random matches, protein pairs were considered crosslinked only if there were at least three unique crosslinked residue pairs between them.

For crosslinked D2-I samples, xiSEARCH version 1.7.6 was used to enable downstream quantitation of crosslinks. The peak lists were searched against the cognate protein sequences and the reversed protein sequences were used as decoys during the search for error estimation. Search results were filtered using xiFDR (version 2.1.5.2). A false discovery rate cut-off was set to 5% at the CMS level, 5% at the crosslinked peptide level and 3% at the residue-pair level. The “Unique PSM” option was disabled and the “boost between” option was enabled. Data from the BS3 crosslinked D2-IWT and D2-I3D were combined for FDR calculation. The same was done for the sulfo-SDA crosslinked D2-IWT and D2-I3D.

MS1-extracted ion-chromatogram-based label-free quantitation of cross-links was carried out using Skyline as previously described^11^. BS3 crosslinked and SDA crosslinked data were quantified separately. In short, the sequences of cross-linked peptides were linearized, and a Skyline input file (in. ssl format) was generated using the xi_Skyline Converter (https://www.rappsilberlab.org/software/xisearch). MS1 signal intensities of each cross-linked peptide were measured by Skyline and normalized between the replicas of each condition. Cross-linked residue pairs were calculated as the sum of the signal intensity of all supporting cross-linked peptides and compared between different D2-I versions. The median normalized log2(D2-I^3D^/D2-I^WT^) ratios were calculated using Perseus.

The MS data of the BS3 crosslinked FA core complex and the sulfo-SDA crosslinked D2-I heterodimers have been previously published^27,45^. The MS raw data were reprocessed in this study as described above.

### Generating the homology model of the chicken FA core complex

We utilized SWISS-Model sequence alignment^51^ to create a homology model of the Fanconi Anemia complex for *Gallus gallus*. The *Gallus gallus* sequences for FANCA, FANCB, streptavidin II tagged FANCC, FANCE, FANCF, FANCG, FANCL, and FAAP100 were provided to the SWISS-Model server as target sequences, and the atomic structural model of the human Fanconi anaemia core complex (PDB|7KZP) was used as the template.

A “single FANCA” model of the *Gallus gallus* FA core complex was generated from the homology model of the complex. The chain A copy of FANCA was retained in the model as its presence is confirmed by the EM density. The other copy of FANCA, chain M, was removed from the homology model using ChimeraX^52^.

### Prediction of the IDRs in the human CCR4-NOT complex

We assign IDRs based on AlphaLink2^53^ predictions and through visual inspection. The predicted lDDT-Cα (pLDDT) has been shown to be a good indicator^54^. We select unfolded regions in the prediction that have low confidence with a cutoff of pLDDT < 30.

### Generating the mutant human CCR4-NOT complex

Point mutants in the CNOT1 subunit were generated using overlapping PCR primers to amplify a codon optimised ORF within a pACEBac1 vector backbone. Individual gel purified PCR fragments were reassembled into a new mutant pACEBac1-CNOT1 vector using NEBuilder® HiFi DNA assembly (NEB) and validated using both Sanger and Nanopore based sequencing. Mutant CNOT1 was then assembled into a single vector containing expression cassettes for all core CCR4-NOT subunits using a modified BigBac protocol. Viruses for co-expression of complexes were prepared as described in the methods for the WT complex.

### Differential scanning fluorimetry

Complex thermostability was assayed by intrinsic fluorescence recorded using a Prometheus NT.48 nano-DSF instrument (NanoTemper technologies), essentially as previously described ^55^. Purified complexes were diluted to 0.25 mg/ml in 20 mM HEPES pH 8, 150 mM NaCl, 0.5 mM TCEP and 2 mM Mg acetate. Data were analysed in ThermControl software (v2.3 NanoTemper) and plotted using Prism (GraphPad).

### In vitro deadenylation assays of the mutant human CCR4-NOT complex

Targeted activity assays were performed as previously described ^56,57^ but with conditions optimised for the human complex . Substrate RNA was synthesized with a 5’ 6-FAM fluorophore and contained a Pumilio response element (PRE) embedded within a 20-mer non-poly(A) region, upstream of a 30-mer poly(A) tail. CCR4-NOT was diluted to 0.5 µM (10x stock) in 20 mM HEPES pH 8, 300 mM NaCl, 2 mM Mg Acetate and 0.5 mM TCEP. Full-length MBP-tagged human PUM1 was diluted to 2.5 µM (10x stock) in 20 mM HEPES pH 8, 150 mM KCl and 0.5 mM TCEP. Assays were performed in 100 µl total volume in a thermally controlled block at 22 °C. RNA at a final concentration of 200 nM was preincubated with 250 nM PUM1 for 15 minutes in deadenylation buffer (20 mM HEPES pH 7.8, 80 mM NaCl, 0.1 mM TCEP, 2 mM MgAc). Reactions were then started by the addition of 10x CCR4-NOT to a final concentration of 50 nM. 7.5 µl samples at the indicated time points were withdrawn and mixed with an equivalent volume of loading dye (95% Formamide, 4 mM EDTA, 0.01 w/v bromophenol blue). Deadenylated species were resolved on denaturing TBE (Tris-borate-EDTA) 20% polyacrylamide (19:1) gels containing 7 M urea and run at 400 V in 1 × TBE running buffer. Gels were visualised using a Amersham Typhoon 5 system (Cytiva) using a 488 nm excitation laser and 525 nm 20 nm bandpass emission filter.

### Data visualization

xiVIEW^58^ was employed to visualize the crosslinking data and generate crosslinking maps in both circle and bar views for figure preparation. The Euclidean distances between the Cα atoms of crosslinked residue pairs in the atomic models of proteins were measured using xiVIEW. ChimeraX^52^ and PyMOL (The PyMOL Molecular Graphics System, version 2.0.7, Schrödinger, LLC) were utilized to display crosslinks within 3D structural models for figure preparation. Additionally, ChimeraX was used to visualize the EM density map of the chicken FA core complex (EMD-10290, data values range from -0.0282 to 0.0628, contour level 0.005393) and to fit the homology model of the complex into the EM density map.

## Extended data

**Extended data Figure 1:**
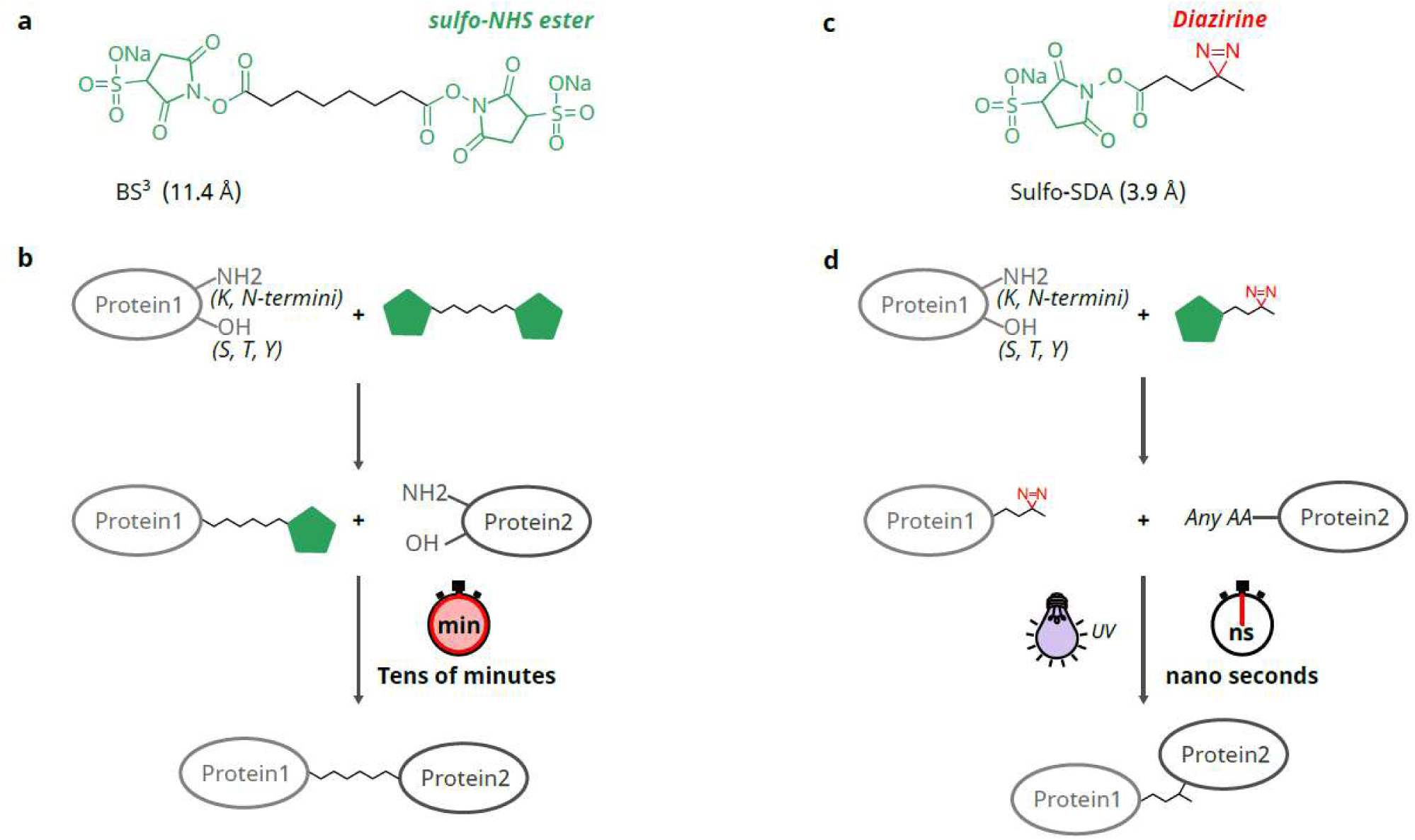
Crosslinking reactions of crosslinkers BS3 and sulfo-SDA. a. The structure of BS3, with the sulfo-NHS ester reactive group highlighted in green. b. Crosslinking reactions of BS3. The green pentagon represents the sulfo-NHS ester. BS3 primarily reacts with the -NH2 groups in lysine side chains and protein N-termini, as well as the -OH groups in threonine, tyrosine, and serine side chains. Once BS3 has reacted with a protein at one end, the NHS ester group on the other end remains reactive for several minutes, allowing the formation of a crosslink when another reactive residue is encountered. c. The structure of sulfo-SDA, with the sulfo-NHS ester highlighted in green and the diazirine highlighted in red. d. Crosslinking Reactions of sulfo-SDA. Sulfo-SDA undergoes a two-step crosslinking reaction. First, the sulfo-NHS ester reacts with proteins identically to BS3. In the second step, the diazirine group on the opposite end is activated by UV light to react with any proximal amino acid (AA), forming a photo-crosslink. The activated diazirine has an extremely brief lifetime of just a few nanoseconds, during which it can either form a crosslink or be hydrolyzed.

**Extended data Figure 2:**
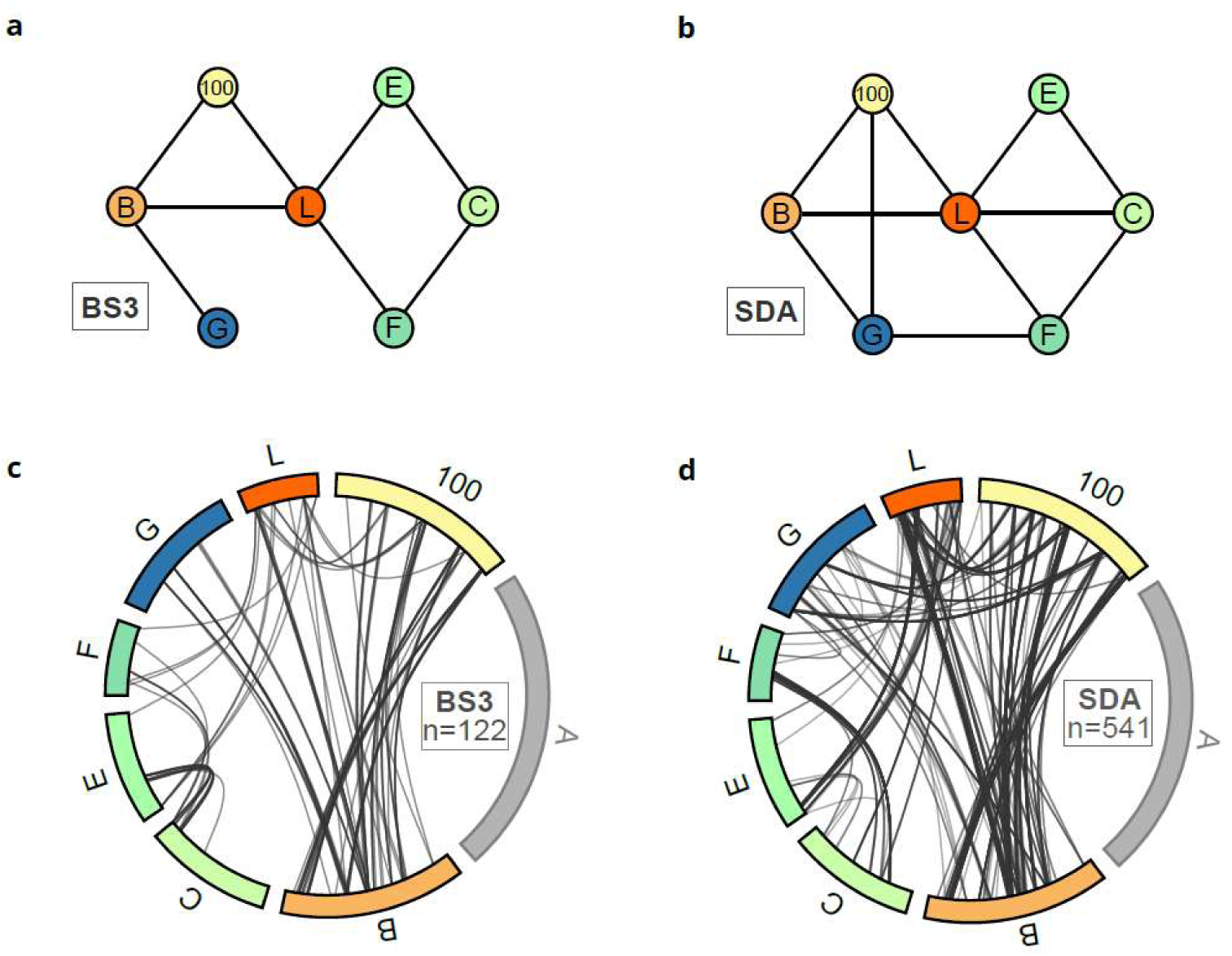
Similarity of BS3 and SDA crosslinks between the subunits of the FA core complex other than FANCA. a. BS3 crosslinks between the non-FANCA subunits in the FA core complex. b. SDA crosslinks between the non-FANCA subunits in the FA core complex. c. A map of 122 BS3 crossliks between the non-FANCA subunits in the FA core complex. d. A map of 541 SDA crossliks between the non-FANCA subunits in the FA core complex.

**Extended data Figure 3.**
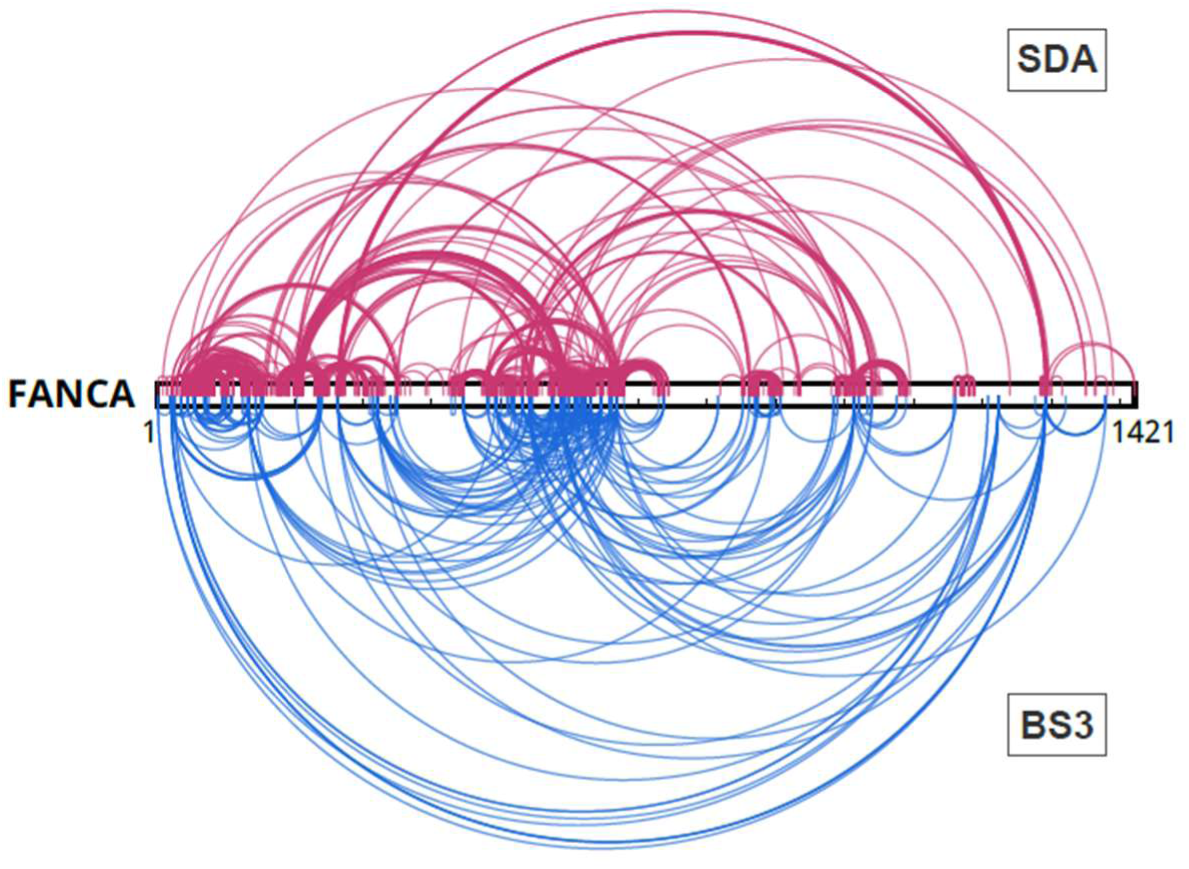
SDA and BS3 self crosslinks of the FANCA subunit. FANCA is shown as a bar. SDA crosslinks are shown in dark pink and BS3 crosslinks are shown in blue.

**Extended data Figure 4:**
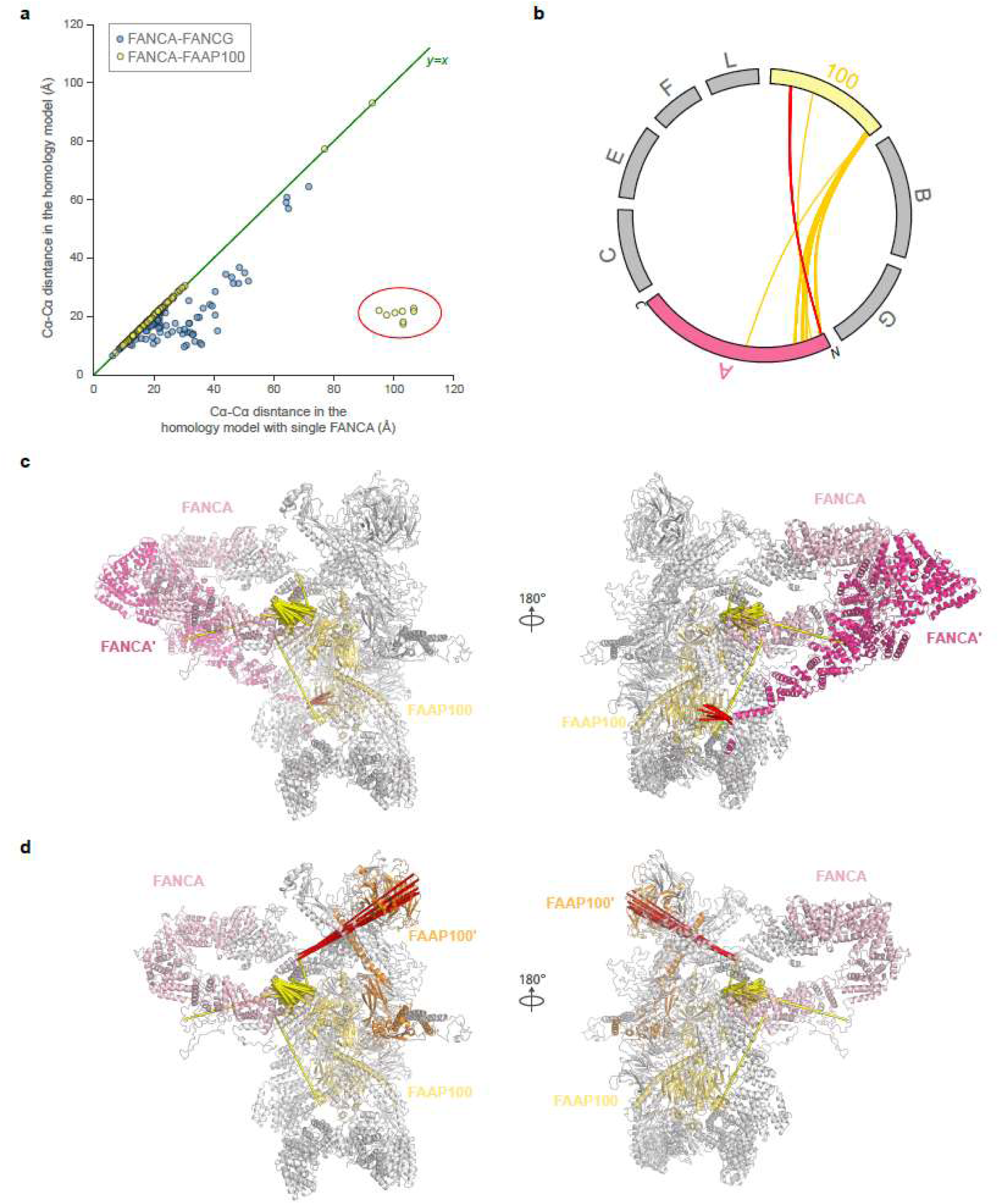
SDA crosslinking data suggest the presence of two copies of FANCA in the chicken FA core complex. a. The Cɑ-Cɑ distances of FANCA-FAAP100 crosslinks (yellow) and FANCA-FANCG crosslinks (blue) measured in the homology model containing two copies of FANCA were plotted against the Cɑ-Cɑ distances of these crosslinks in the model of the FA core complex containing a single copy of FANCA. Some other subunits are present in multiple copies in the structural models, and in those cases, residue pairs that have the shortest Cɑ-Cɑ distances were used. Eight FANCA-FAAP100 crosslinks show agreement only with the homology model with two FANCAs, highlighted in the red oval. b. A map of 91 FANCA-FAAP100 crosslinks. The eight crosslinks that are highlighted in panel A are shown in red and are between the N-terminal region of FANCA and the C-terminal region of FAAP100. c. The 91 FANCA-FAAP100 crosslinks are displayed in the homology model of the FA core complex. The two copies of FANCA are coloured in light pink (marked as A) and in hot pink (marked as A’). One of the two copies of FAAP-100 that are crosslinked to FANCA/FANCA’ is coloured in yellow and marked as 100. Crosslinks are shown as yellow strokes, and the eight crosslinks highlighted in panel A are coloured in red. d. The 91 FANCA-FAAP100 crosslinks are displayed in the model of the FA core complex with only one FANCA. FANCA is coloured light pink. The two copies of FAAP100 are coloured in yellow (marked as 100) and in light orange (marked as 100’)

**Extended data Figure 5:**
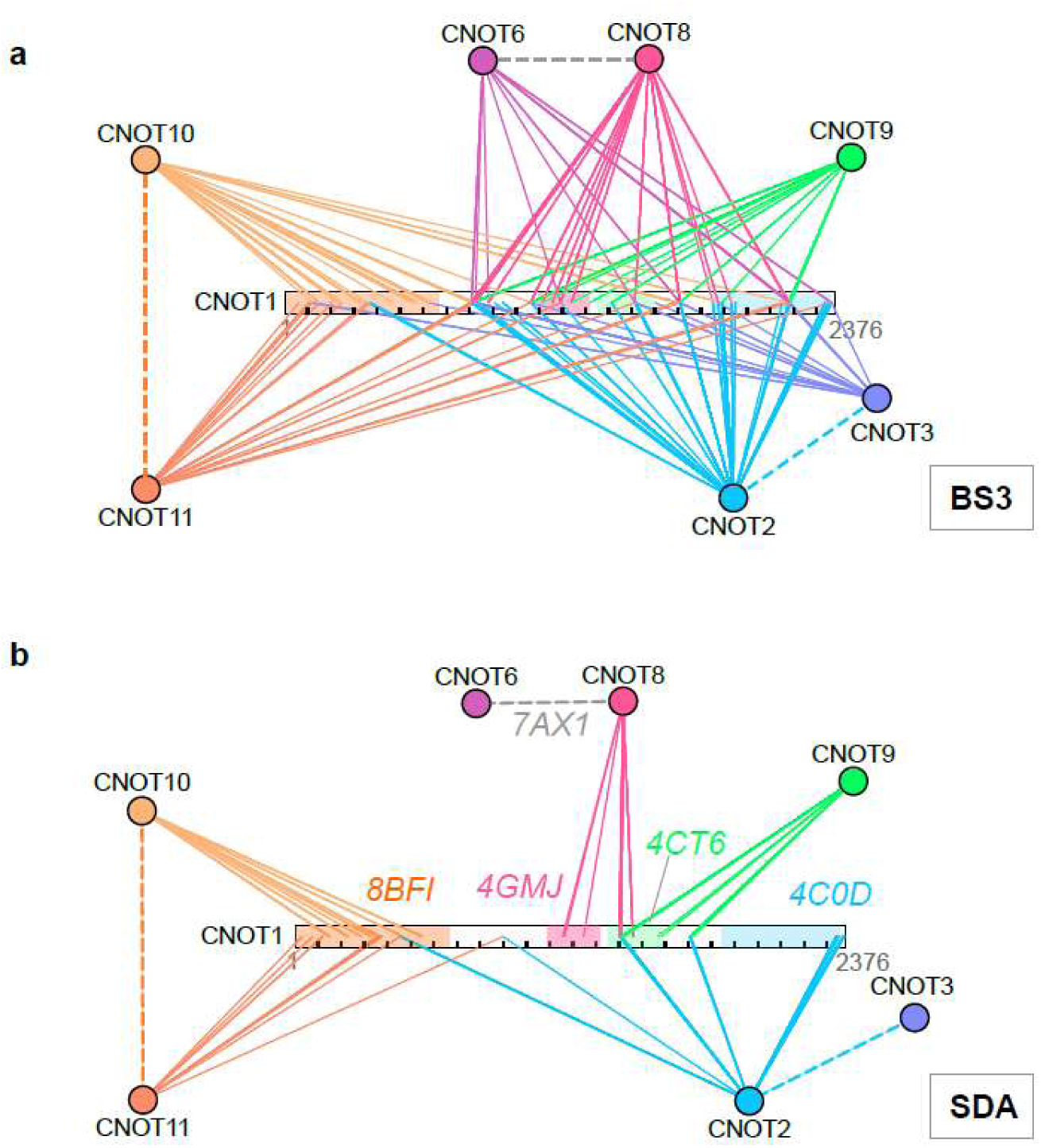
Crosslinks between CCR4NOT subunits and the scaffold subunit CNOT1. 289 BS3 crosslinks (panel a) and 173 SDA crosslinks (panel b) between the scaffold subunit CNOT1 and other subunits of the human CCR4-NOT complex. CNOT1 is shown as a bar and other subunits are shown as colored circles. Crosslinks are shown as solid lines and coloured according to the colour of the non-CNOT1 subunits. The dashed lines indicate known interactions between the subunits. CNOT1 regions that are known to interact with other CCR4-NOT subunits in four functional modules are highlighted in coloured shades: CNOT1^1–685^ which interacts with CNOT10-CNOT11 (orange), CNOT1^1093–1317^ which interacts with CNOT6-CNOT8 (pink), CNOT1^1356–1588^ which binds CNOT9 (green) and CNOT1^1847–2361^ which interacts with CNOT2-CNOT3 (blue). Corresponding crystal structures that revealed the interactions between subunits are indicated.

**Extended data Figure 6:**
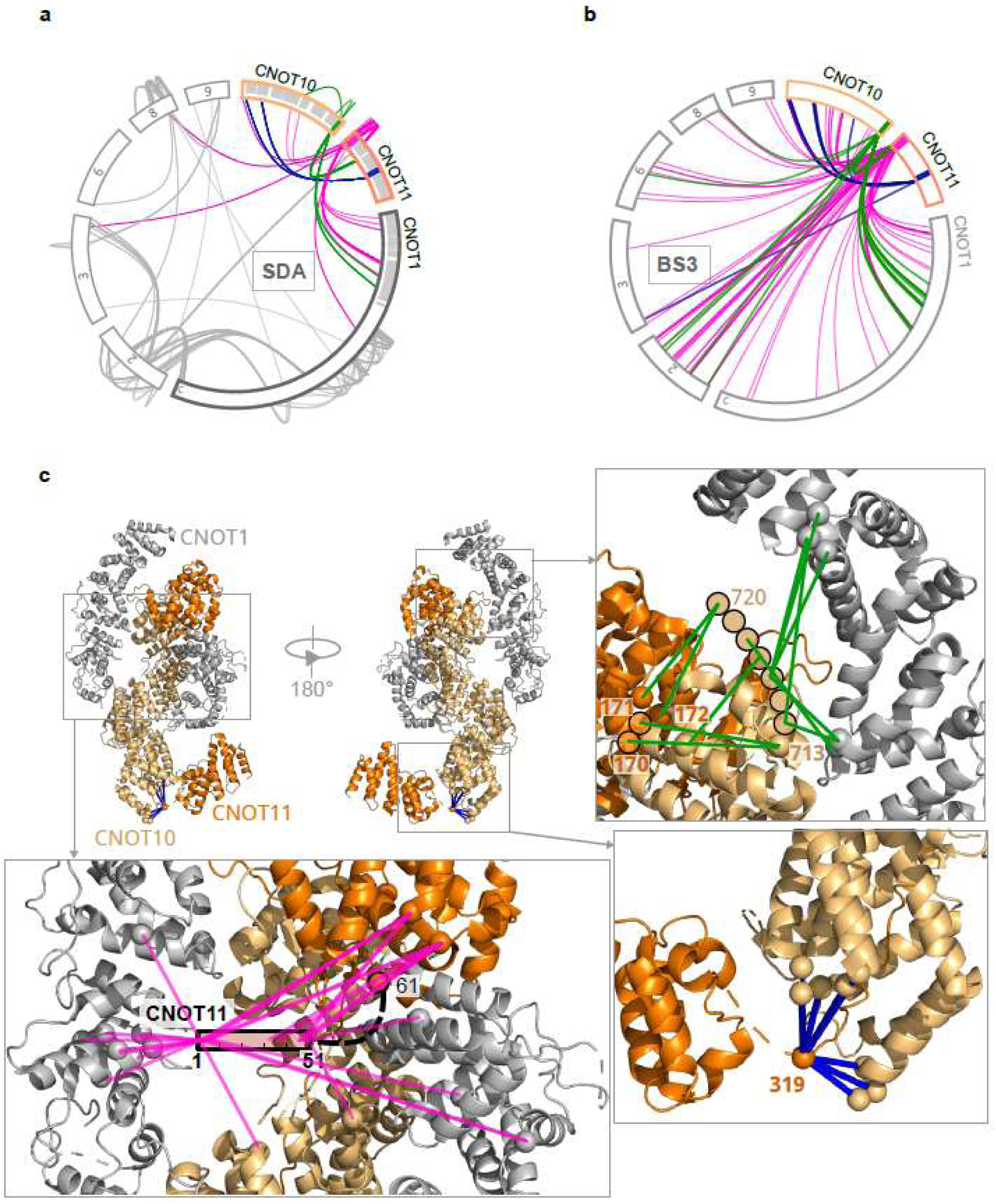
IDR interactions in the CCR4-NOT complex. a. SDA crosslinks from the C-terminal IDR in CNOT10 (707-744, highlighted in green), and IDRs at the N-terminal (1-60 highlighted in pink) and central regions of CNOT11 (285-320 highlighted in blue). These crosslinks are highlighted on a map of all SDA IDR crosslinks observed in the human CCR4-NOT complex. The crosslinks are coloured based on which IDR they are linked to while all other crosslinks are shown in gray. The gray half shades indicate residues that are present in the crystal structure of the CNOT1-CNOT10-CNOT11 module (PDB 8BFI) b. BS3 crosslinks from the three IDRs described in panel a. The crosslinks are coloured based on which IDR they are linked to. c. The SDA crosslinks from the three IDRs described in panel a visualised on the crystal structure of the CNOT1-CNOT10-CNOT11 module (PDB 8BFI). Three regions from the crystal structure are marked and each displayed in an amplified view. On the upper right view, crosslinks from the C-terminal IDR in CNOT10 are displayed as green strokes. Crosslinked residues in the crystal structure are shown as spheres while crosslinked residues in CNOT10^714–720^, CNOT11^170–171^ which are absent in the crystal structure, are shown as spheres with black outlines, extending from residues CNOT10^713^ and CNOT11^172^ in the crystal structure. In the lower right view, crosslinks from the IDR at the centre of CNOT11 are displayed as blue strokes, while crosslinked residues in the crystal structure are shown as spheres. In the lower-left view, CNOT11^1–51^ of the N-terminal IDR is shown as a bar connecting to CNOT11^61^, the most N-terminal CNOT11 residue in the crystal structure. Crosslinks from this IDR to the residues in the crystal structure are shown as pink strokes. The crosslinked residues in the crystal structure are shown as spheres.

**Extended data Figure 7:**
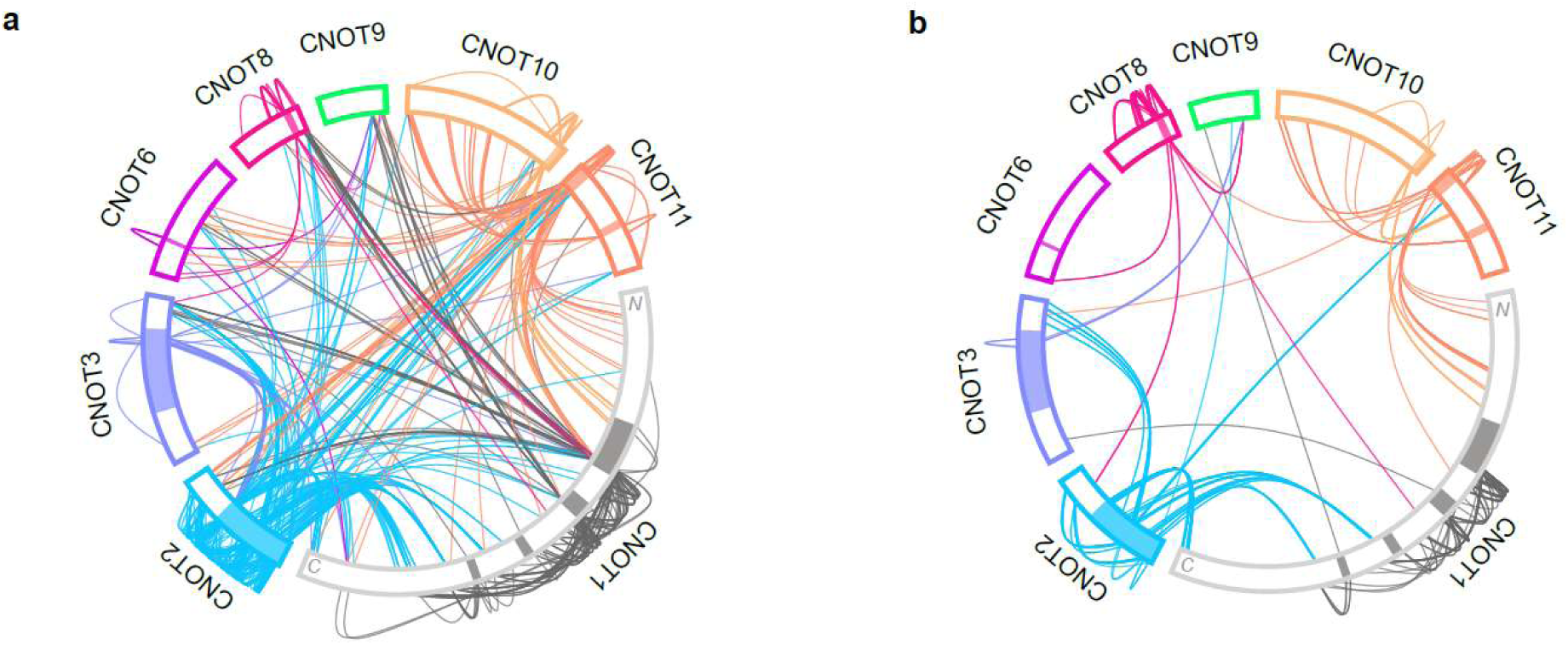
Crosslinks from IDRs in the human CCR4-NOT complex. Crosslink map of 631 BS3 crosslink (panel a) and 321 SDA crosslinks (panel b) from the IDRs in the human CCR4-NOT complex. IDRs are highlighted with gray shade. The crosslinks are coloured based on the protein colour of the IDR it belongs to.

**Extended data Figure 8:**
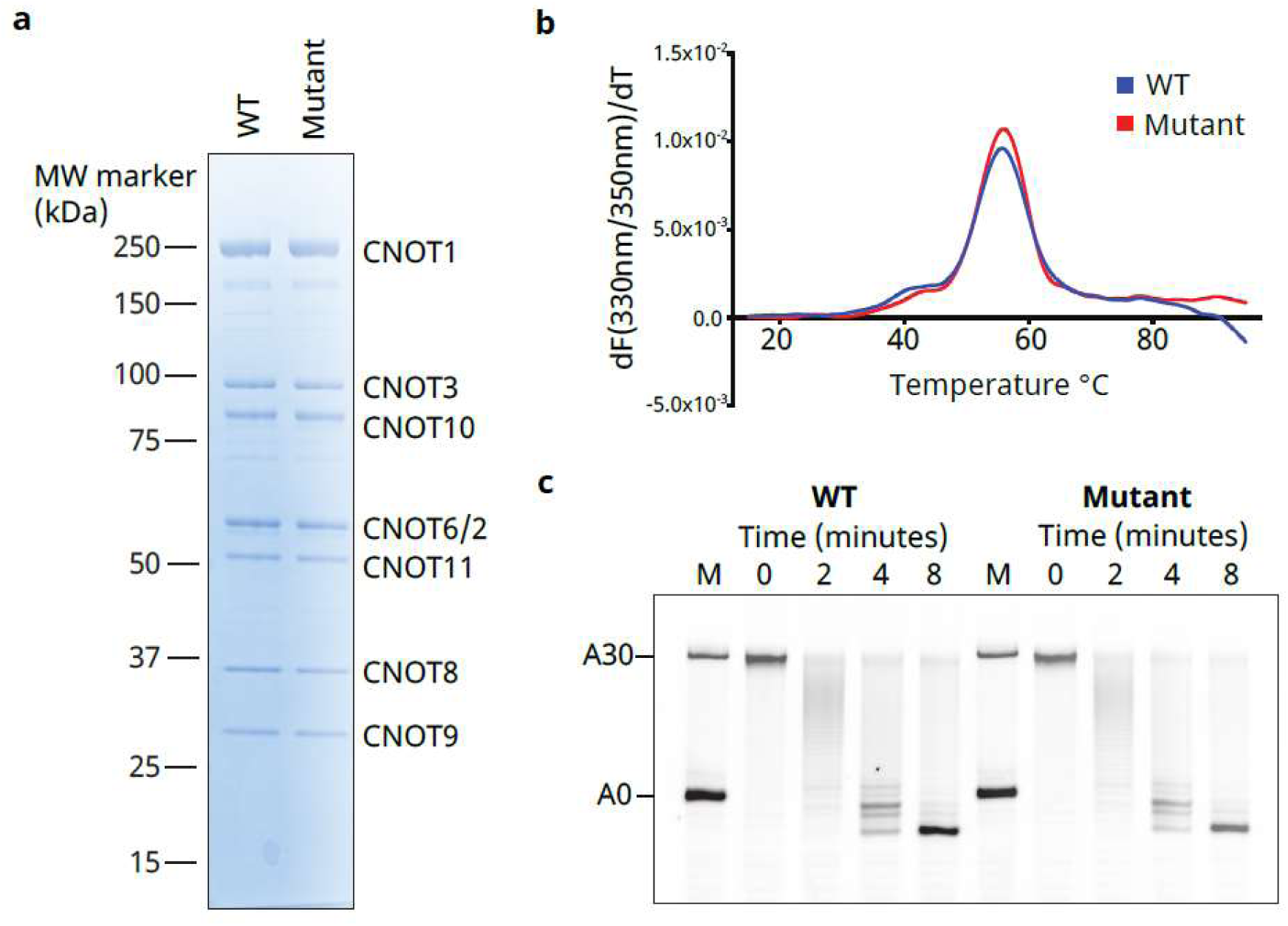
Mutation of CNOT1 IDR-binding interface. a. Coomassie stained 4-12% gradient SDS-PAGE analysis of purified WT and mutant recombinant human CCR4-NOT complexes. The mutant complex carries the following mutations in the CNOT1 subunit: L1395K, Q1430A, L1432T, V1433E, H1434S. All subunits are present in unit stoichiometry, with fainter bands corresponding to proteolytic fragments. b. Differential scanning fluorimetry of WT (blue) and mutant (red) complexes. The intrinsic fluorescence of complexes (0.25 mg/ml) at 330 and 350 nm was measured as a function of temperature using a Prometheus instrument and the first differential was plotted to determine the unfolding transition. Both WT and mutant complexes show a similar unfolding pattern and thermostability. c. *In vitro* deadenylation assays of recombinant WT and mutant complexes in the presence of the RNA-binding protein PUM1 (interacting with the NOT module of the CCR4-NOT complex). The substrate is a fluorescently-labelled synthetic RNA containing an upstream PUF-recognition element (PRE) followed by a 30-nt poly(A) tail. The reaction mixture was resolved using denaturing PAGE. The sizes of RNA with an A30 tail and no poly(A) tail is indicated.

**Extended data Figure 9:**
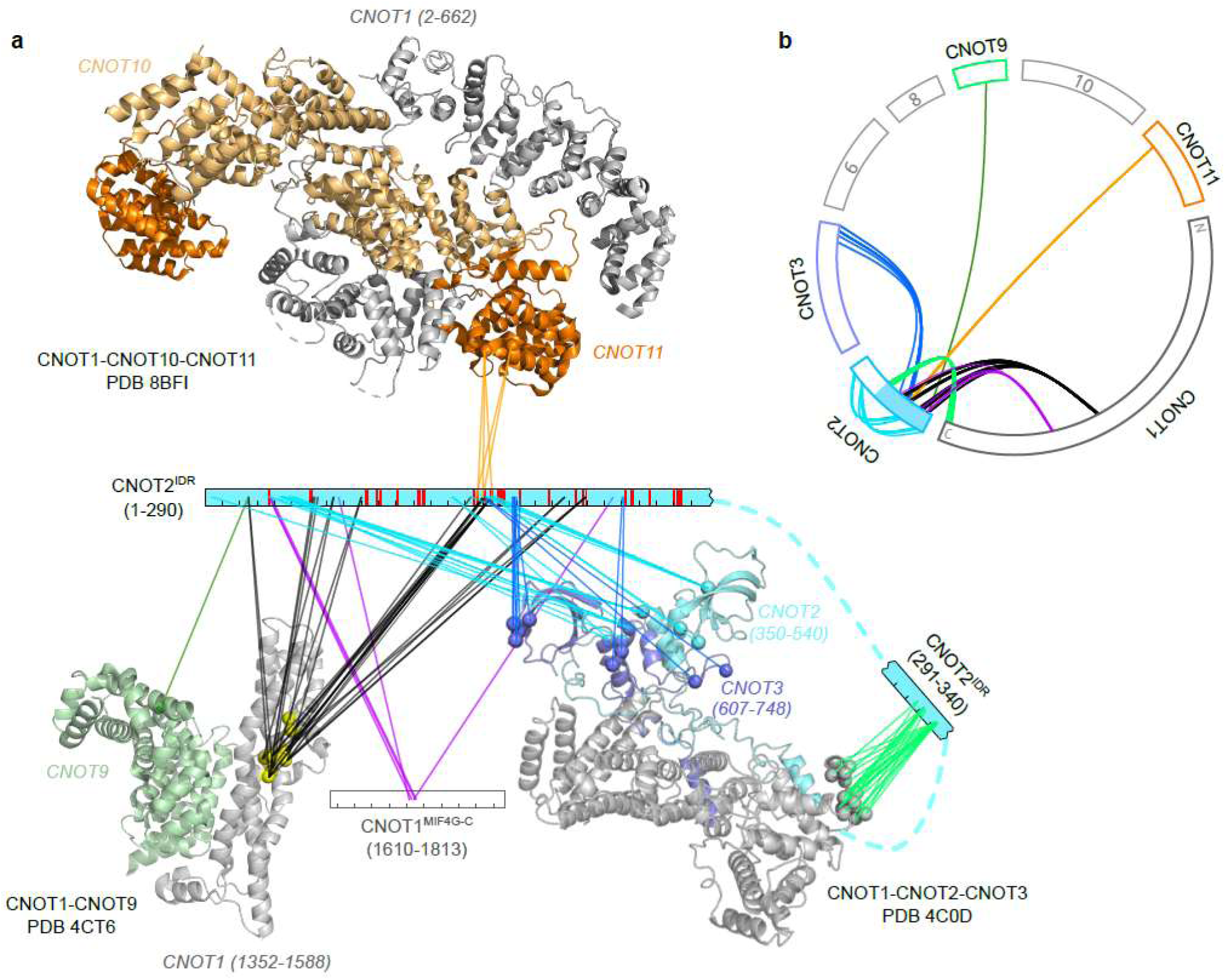
A crosslink network of CNOT2^IDR^. a. A network of crosslinks from the CNOT2^IDR^ to the structured regions in the other subunits in the human CCR4-NOT complex. The CNOT2^IDR^ is shown as a blue bar separated into two segments (1-290 and 291-340). Crosslinks from the CNOT2^IDR^ are depicted as strokes to the linked residues in the crystal structures of the CNOT1-CNOT2-CNOT3 module (PDB 4C0D), the CNOT1-CNOT9 module (PDB 4CT6), and the CNOT1-CNOT10-CNOT11 module (PDB 8BFI), or the white bar that represents the CNOT1^MIF4G-C^ domain. The crosslinks are colored according to the structured regions they link to. The crosslinked residues in the crystal structures are indicated as spheres. The known phosphorylation sites in the CNOT2^IDR^ are indicated in red. b. The crosslinks displayed in panel A are shown in a crosslink map.

**Extended data Figure 10:**
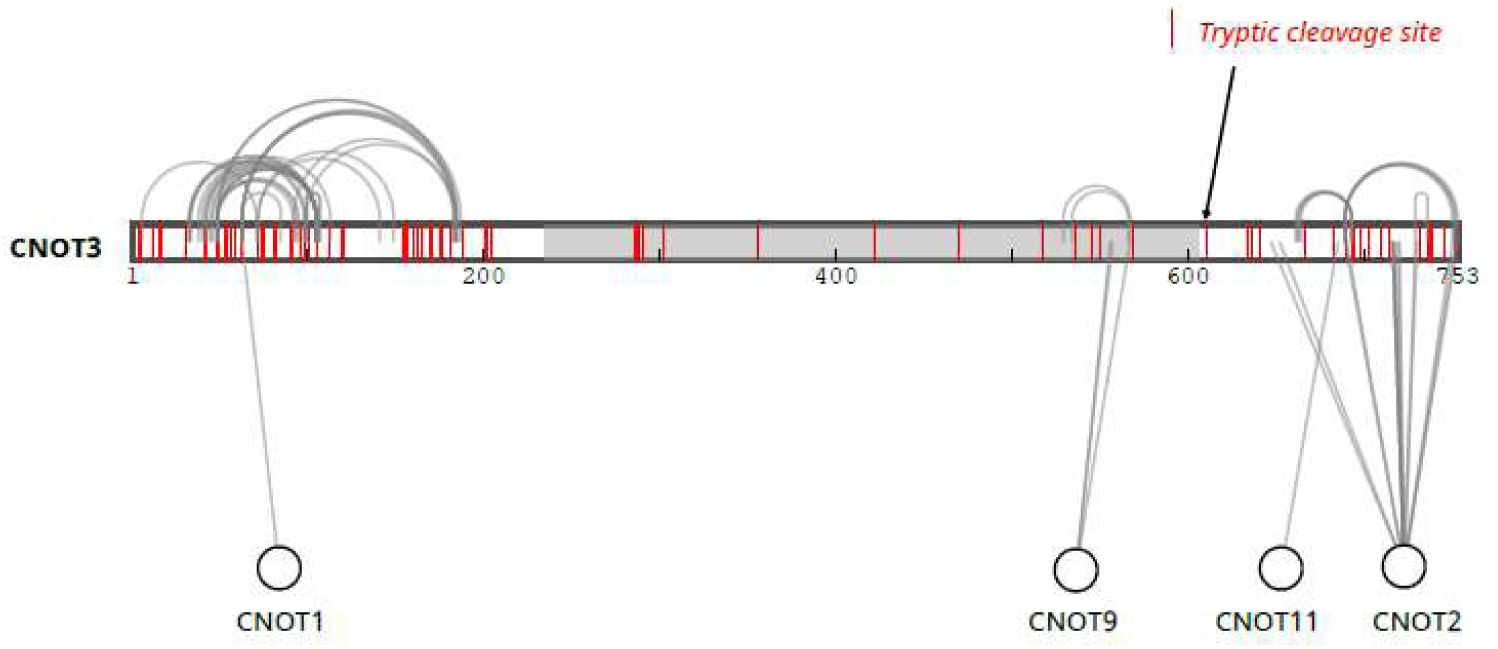
Lack of trypsin cleavage sites impaired the detection of crosslinks in the middle region of CNOT3. CNOT3 is represented as a bar, with tryptic cleavage sites indicated by red strokes on the bar. The IDR region in CNOT3 is shaded. SDA crosslinks of CNOT3 are depicted as archs (self-links) and solid lines (heterometric-links). Four proteins that crosslinked to CNOT3 are represented by circles.

## Notes

### Competing Interest Statement

The authors have declared no competing interest.

